# The concerted action of SEPT9 and EPLIN modulates the adhesion and migration of human fibroblasts

**DOI:** 10.1101/2022.08.08.503265

**Authors:** Matthias Hecht, Pia Marrhofer, Nils Johnsson, Thomas Gronemeyer

**Affiliations:** Institute of Molecular Genetics and Cell Biology, James Franck Ring N27, Ulm University, 89081 Ulm, Germany

## Abstract

Septins are cytoskeletal proteins that participate in cell adhesion, migration and polarity establishment. The septin subunit SEPT9 directly interacts with the single LIM domain of EPLIN, an actin bundling protein. Using a human SEPT9 KO fibroblast cell line we show that cell adhesion and migration are regulated by the interplay between both proteins. The low motility of SEPT9-depleted cells could be partly rescued by increased levels of EPLIN. The normal organization of actin-related filopodia and stress fibers was directly dependent on the expression level of SEPT9 and EPLIN. Increased levels of SEPT9 and EPLIN enhanced the size of focal adhesions in cell protrusions, correlating with a stabilization of actin bundles, whereas decreased levels have the opposite effect. Our work thus establishes the interaction between SEPT9 and Eplin as important link between the septin and the actin cytoskeleton that influences cell adhesion and motility.

## Introduction

Septins were discovered in the budding yeast in the 1970s and were subsequently shown to form a novel cytoskeletal system (Longtine et al., 1996). Unlike actin filaments and microtubules, septins assemble into non-polar filaments. The mammalian genome encodes thirteen different septins (SEPT1-SEPT12, SEPT14) (Shuman and Momany, 2022). The basic septin building block in mammalian cells is a hetero-octamer composed of the SEPT2, SEPT6, SEPT7, and SEPT9 at a stoichiometry of 2:2:2:2 (2-7-6-9-9-6-7-2) (Mendonça et al., 2019). These building blocks are capable of polymerizing by end-to-end and lateral joining into higher ordered structures such as rings, filaments, and gauzes. Initially labeled as passive scaffold proteins, it became more and more evident that septins are actively involved in many intracellular processes. They regulate vesicle transport and fusion, chromosome alignment and segregation, and cytokinesis (Surka et al., 2002; Bowen et al., 2011; Fuechtbauer et al., 2011; Tokhtaeva et al., 2015; Estey et al., 2013). In addition, septins cross-link and bend actin filaments into functional structures such as contractile rings during cytokinesis, or stress fibers in filopodia and lamellipodia (Mavrakis et al., 2014; Dolat et al., 2014). Although actin and septin networks only partially overlap, they are structurally interdependent as disruption of actin alters the organization of septin networks and vice versa.

Cell migration is mainly accomplished by a turnover of actomyosin stress fibers and focal adhesions. Septins are known to colocalize with stress fibers, enriched in the leading lamella and in radial stress fibers anchored to focal adhesions (Dolat et al., 2014). Several studies connected the septins to mechanotransduction and therefore to the maintenance of front-rear polarity in migratory cells (Lam and Calvo, 2019; Simi et al., 2018; Calvo et al., 2015; Dolat et al., 2014). Accordingly, the depletion of septins leads to a loss of the front-rear polarity axis of migrating cells (Shindo et al., 2018). Loss of cell-cell adhesion and alteration of cell polarity is also often observed in tumors of epithelial origin (Pal et al., 2018). These changes correlate with the altered SEPT9 expression levels in diverse types of epithelial cancer including prostate, breast and colon cancer (Connolly et al., 2011; Song and Li, 2015; Gilad et al., 2015; Verdier-Pinard et al., 2017). Despite these many supporting observations, it is still elusive how SEPT9 is mechanistically linked to the adhesion and migration machinery.

The LIM domain containing protein EPLIN (epithelial protein lost in neoplasm) is a modulator of cellular architecture and the actin cytoskeleton (Maul et al., 2003; Zhang et al., 2011) and was recently identified as a binding partner of SEPT9 (Hecht et al., 2019). EPLIN crosslinks actin filaments to bundles and thereby inhibits the Arp2/3-mediated depolymerization and branching of filaments (Maul and Chang, 1999). Whereas the binding of EPLIN to the pointed end of actin filaments decreases depolymerization, the nucleation by Arp2/3 is inhibited and thus the dynamic turnover of the actin cytoskeleton is reduced. Low intracellular EPLIN levels correlate with an enhanced cancer cell invasion that is partially induced by a lack or loss of the tumor suppressor p53 (Ohashi et al., 2017). Epithelial-mesenchymal transition (EMT), characterized by a loss of apico-basal polarity and abnormal alterations of cell shape and organization, is promoted by a reduction or loss of EPLIN (Zhang et al., 2011). A physical interaction of EPLIN with the cadherin-catenin complex was shown to be required for the formation of adherens junctions (Abe and Takeichi, 2008). Furthermore, the overall maintenance of cell polarity in epithelial cells depends on the formation of a large protein complex consisting of E-cadherin, β-catenin, α-catenin, EPLIN and F-actin (Chervin-Pétinot et al., 2012). EPLIN responds to mechanical forces. As the LIM domain of EPLIN does not bind to actin, its mode of mechanosensing must differ from the LIM-domain dependent mechanisms of the members of the paxillin and zyxin family (Taguchi et al., 2011; Gulino-Debrac, 2013).

## Results

### SEPT9 interacts with the LIM domain of EPLIN

The epithelial protein lost in neoplasm (EPLIN), the product of the LIMA1 gene, is a regulator of cell-cell/cell-surface adhesion and proliferation and was identified by us as a novel SEPT9 interaction partner (Hecht et al., 2019). EPLIN and SEPT9 co-locate along cytosolic septin fibers but also at the protrusion tip of motile cells and at the cleavage furrow of dividing cells (Fig.1A, B and Video 1) in 1306 fibroblasts. We expressed both proteins recombinantly in *E. coli* and could show that purified SEPT9 interacts in a zinc dependent manner with immobilized GST-EPLIN (Fig.1C). EPLIN is a largely unstructured protein with one central LIM domain. The LIM domain contains a double zinc finger motif that is responsible for the correct folding of the domain (Michelsen et al., 1993). The zinc-dependency of the interaction thus strongly supports a contribution of the LIM domain in binding to SEPT9. In addition, the interaction of EPLIN with SEPT2 was previously shown by Co-IP form HeLa cell lysates (Chircop et al., 2009). As SEPT2 and SEPT9 only share the central G domain as their common feature, we tested whether the G-domain of SEPT9 directly binds to the LIM domain of EPLIN. Indeed, the SEPT9_295-567_ construct comprising only the G-domain suffices to bind the isolated LIM domain in a zinc dependent manner (Fig. 1D).

**Figure 1.**
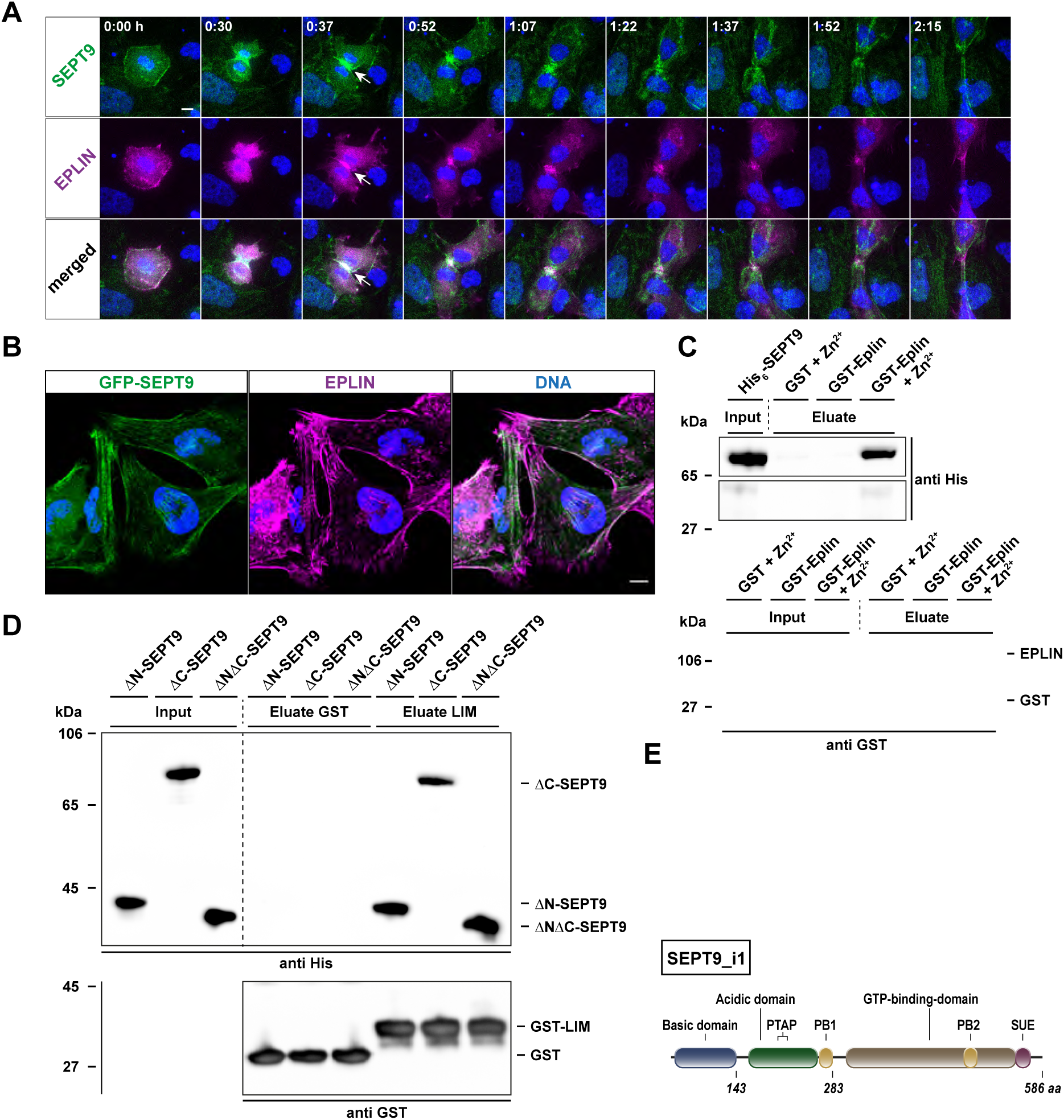
SEPT9 colocalizes and interacts with EPLIN. **(A)** GFP-SEPT9 and mRuby2-EPLIN colocalize at the cleavage furrow in dividing 1306 cells (white arrows). Confocal images were taken every 7.5 min and multiple Z-stacks combined to an average projection (scale bar = 10 µm). **(B)** Immunfluorescence of EPLIN in GFP-SEPT9 expressing cells showing colocalization along septin fibers and at the protrusion of cells (scale bar = 10 µm). **(C)** Pulldown showing the Zn^2+^ dependent interaction of purified His_6_-SEPT9 with GST alone and GST-EPLIN. A protein concentration of 1.5 µM SEPT9 and 0.1 mM Zn^2+^ were applied. **(D)** Purified, recombinantly expressed His_6_-tagged SEPT9 fragments (ΔN-SEPT9 (aa 295-586), ΔC-SEPT9 (aa 1-567), ΔNΔC-SEPT9 (aa 295-567)) were tested for binding to GST alone and GST-EPLIN_LIM_ in a pull-down assay. The isolated G-domain in the ΔNΔC-SEPT9 construct was sufficient to mediate the interaction with the LIM domain. All assays were performed in presence of 0.1 mM Zn^2+^ with a protein concentration of 1.5 µM. **(E)** Scheme of the domain structures of EPLIN and SEPT9i1.

### The expression level of SEPT9 correlates with the migratory properties of fibroblasts

Increased levels of SEPT9 were shown to enhance the motility of murine cells and of human cancer cell lines (Füchtbauer et al., 2011; Farrugia et al., 2020; Marcus et al., 2019). We generated a 1306 cell line stably overexpressing GFP-SEPT9 to study the effect of SEPT9 on cell motility. Constitutive overexpression led to an increase in the amount of SEPT9 by approximately 30% compared to WT cells (Suppl. Fig. S1A). To study the effect of the loss of SEPT9, we generated a CRISPR/Cas9 mediated knock out (KO) of exons 4-6 in 1306 fibroblasts resulting in an interruption of the SEPT9 ORF. The resulting cell line lacked detectable levels of SEPT9 as evaluated by Western blotting (Suppl. Fig. S2). Cell motility of 1306 wildtype fibroblasts, GFP-SEPT9 overexpressing cells (OE) and two clones of our SEPT9 KO cell line were compared by time-lapse microscopy combined with automated, artificial intelligence assisted single cell tracking. We used the log(MSD) (mean square displacement) over time as a measure for cell mobility. The log(MSD) of SEPT9 OE cells was significantly above the WT level whereas the log(MSD) of SEPT9 KO cells was significantly lower (Fig. 2A, B). This indicates that SEPT9 expression levels in 1306 fibroblasts positively correlate with cell motility. An analysis of the global translocation directionality confirmed that in absence of chemoattractants all investigated cell lines moved randomly (Fig. 2C). The velocity of SEPT9 OE cells was significantly enhanced whereas the SEPT9 KO decreased the velocity of the cells below WT level (Fig. 2D).

**Figure 2.**
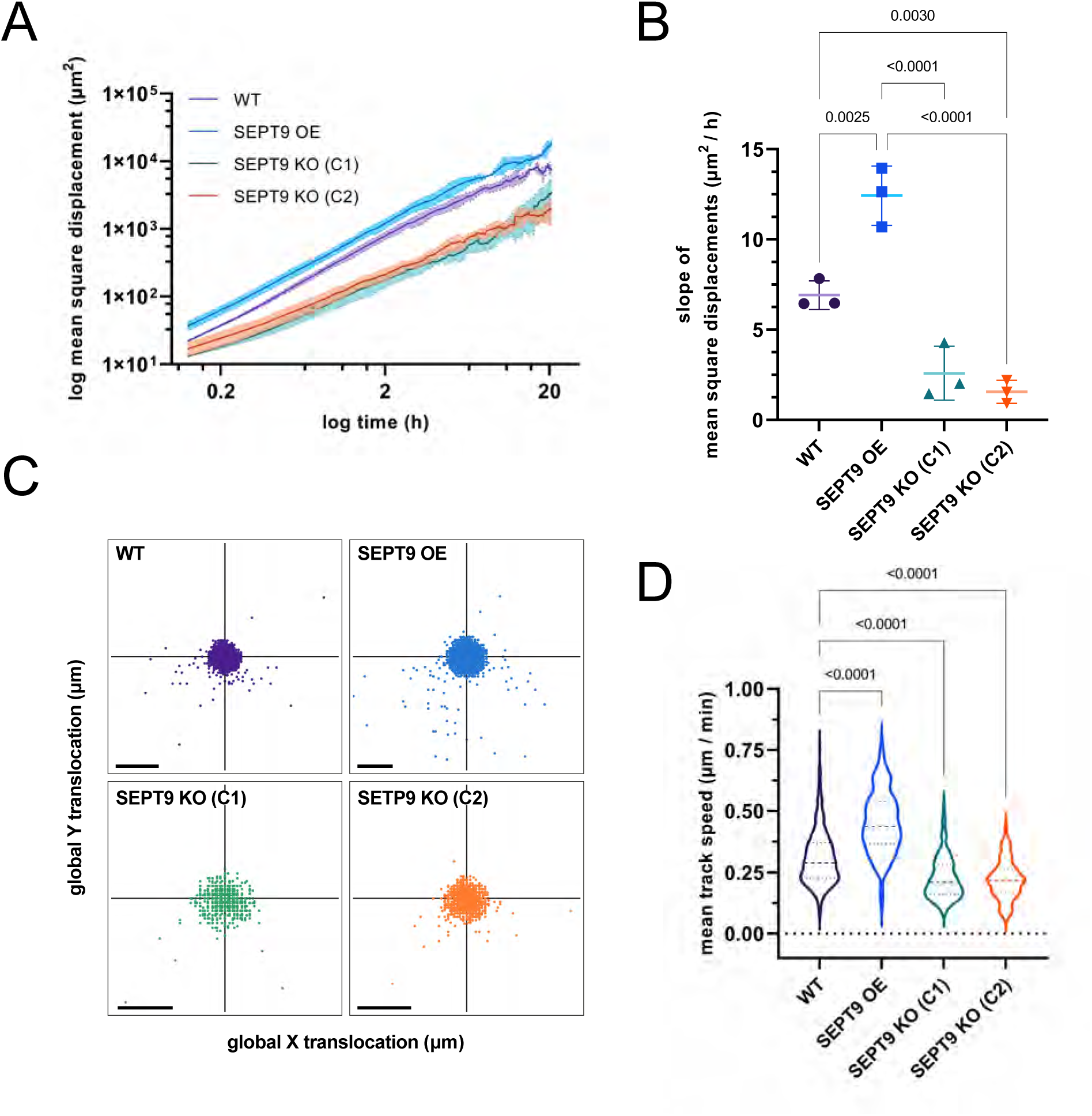
SEPT9 overexpression increases cell motility. **(A)** The mean square displacement (MSD) of 1306 cells changes increases or decreases upon up- or downregulation of SEPT9, respectively. Plotted are the log mean MSD ± range values from three independent experiments. **(B)** The slopes of MSDs differ significantly between SEPT9 KO, WT and SETP9 OE cells. The analysis was performed in triplicate with n = 152 cells. **(C)** The highest density of global step directions per cell line was the center, proving a random movement in chemoattractant-free environment. **(D)** The velocity of cell movement was significantly enhanced upon SEPT9 OE and significantly reduced upon SEPT9 KO (n ≥ 89 cells).

To investigate the effects of the SEPT9 interactor EPLIN on cell migration, we constructed a cell line stably overexpressing GFP-EPLIN. Constitutive EPLIN overexpression led to an increase of the amount of intracellular EPLIN by approximately 50 % compared to WT cells (Suppl. Fig. S1B), whereas downregulation via siRNA knock down (KD cells) reduced the expression level below 10 % in comparison to WT cells and cells transfected with a non-targeting siRNA (Suppl. Fig. S1C). We compared the overall morphology of these and our SEPT9 OE and KO cell lines to wildtype fibroblasts. SEPT9 OE led to an expanded cell morphology while SEPT9 KO led to a constricted morphology. EPLIN OE had no significant impact on the cell morphology while EPLIN KD cells were rounded, resembling a fried egg (Suppl. Fig. S3).

The influence of SEPT9 and EPLIN on cell migration was investigated by a Boyden chamber assay. SEPT9 OE enhanced (1.4-fold increase) the migrative potential of 1306 cells compared to WT cells (Fig. 3A). The KO of SEPT9 stopped the migration of the cells nearly completely (8-fold decrease). Overexpression of EPLIN had only mild effects (1.15-fold increase) on cell migration (Fig. 3B). However, EPLIN KD cells showed a 3-fold higher migration rate as WT cells, or cells expressing a respective control siRNA.

**Figure 3.**
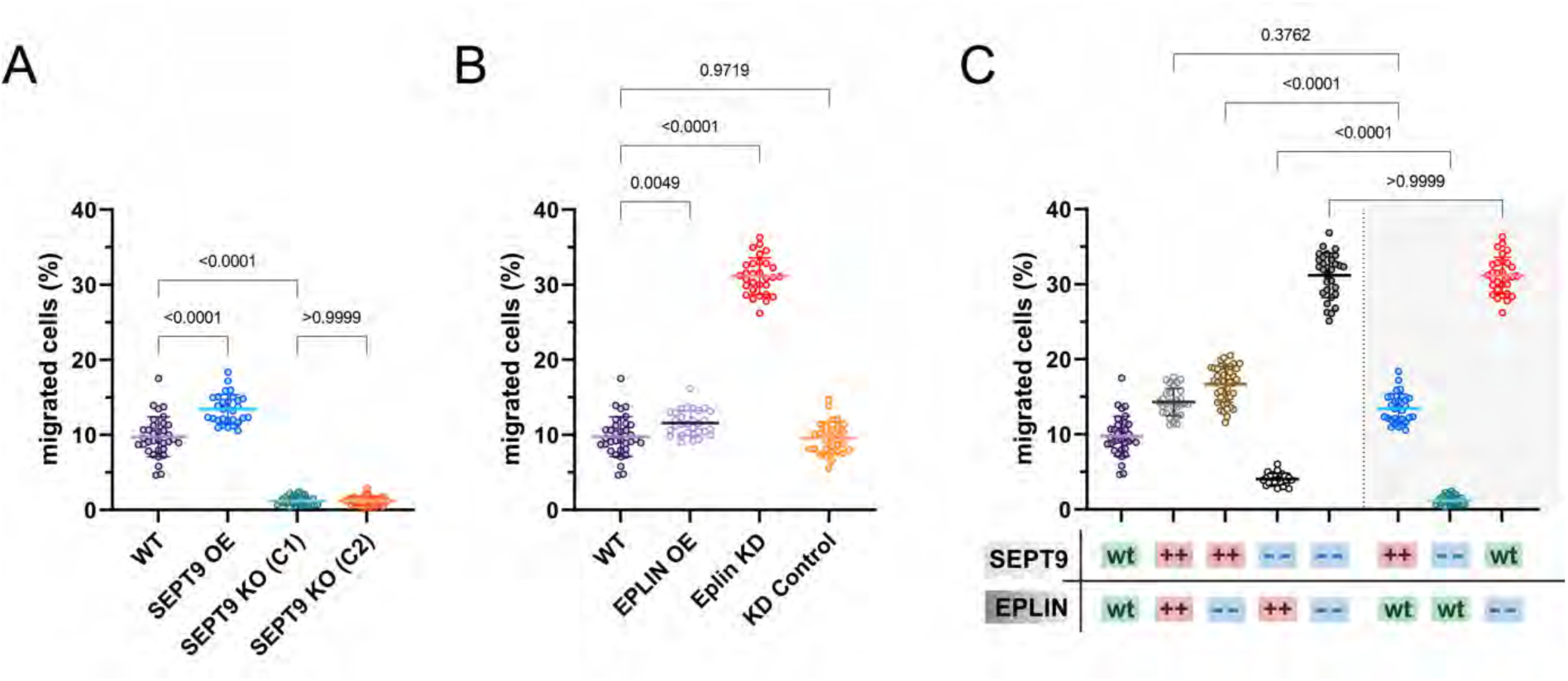
SEPT9 and EPLIN co-regulate cellular migration. **(A)** SEPT9 OE migrate faster than wildtype cells whereas SEPT9 KO cells do barely migrate at all. **(B)** EPLIN KD increases cell migration significantly. **(C)** Cell migration upon simultaneous up- or downregulation of SEPT9 and EPLIN. EPLIN KD can restore the migratory potential of SEPT9 KO cells. Significance values were calculated by one-way ANOVA followed by Šidák’s multiple comparison test from three independent experiments. All quantitative data are depicted as means ± SD.

To investigate the interdependency of the SEPT9 and EPLIN interaction on cell migration, we repeated the Boyden chamber assay with cells where both proteins were simultaneously up- or down-regulated (Fig. 3C). The OE of EPLIN in SEPT9 KO cells resulted only in a minor increase of the migration rate (4 % migrated cells, compared to 1 % in SEPT9 KO). Compared to WT cells (9.7 %), the already enhanced cell migration of SEPT9 OE cells (13.4 %) could not be significantly increased by additional OE of EPLIN (14.3 %). However, the high migration rate of cells with a KD of EPLIN, (31.1 %) was significantly reduced by the simultaneous SEPT9 OE (16.6 %) but was not affected by SEPT9 KO (31.2 %). Taken together, the Boyden chamber assays revealed opposite effects at low levels of SEPT9 (reduced motility) or EPLIN (enhanced motility). High levels of SEPT9 could, in contrast, promote cell migration whereas an OE of EPLIN had no significant impact.

We next asked whether other cellular processes are also co-regulated by SEPT9 and EPLIN. Migration involves breakdown and re-establishment of adhesion to the attachment matrix (Gardel et al., 2010). Malignant transformation is also correlated with the loss of cellular adhesion (Janiszewska et al., 2020). As both proteins were associated with the metastasizing character of various cell lines (Zeng et al., 2019; Sun et al., 2015; Verdier-Pinard et al., 2017; Farrugia et al., 2020; Zhang et al., 2011), we monitored cell-surface adhesion under varying SEPT9 or EPLIN expression levels. Upon seeding of detached cells, the progress of reattachment and cell spreading was documented every 15-30 min by light microscopy. The fraction of attached cells and the degree of spreading was classified in five gradations ranging from 0 to 100 %. First the initiation of attachment for each cell line was determined (Fig. 4A). If at least 25 % of all cells attached to the surface, the elapsed time was counted as “initiation of attachment”. EPLIN OE and EPLIN KD had no influence on the cell-surface interaction, documented by an initial attachment time of 12.5 min almost identical to WT cells. SEPT9 showed strong effects upon OE as the time required for initial attachment was reduced to 7.5 min. In contrast, SEPT9 KO cells exhibited a drastically prolonged attachment time of 52.5 min. Although variation of EPLIN levels alone did not show any effects, the OE of EPLIN in SEPT9 KO cells could partially restore the WT-like attachment initiation to 20 min. A similar effect of EPLIN overexpression was observed for the progress of adhesion of the SEPT9 KO cell line (Fig. 4B, C).

**Figure 4.**
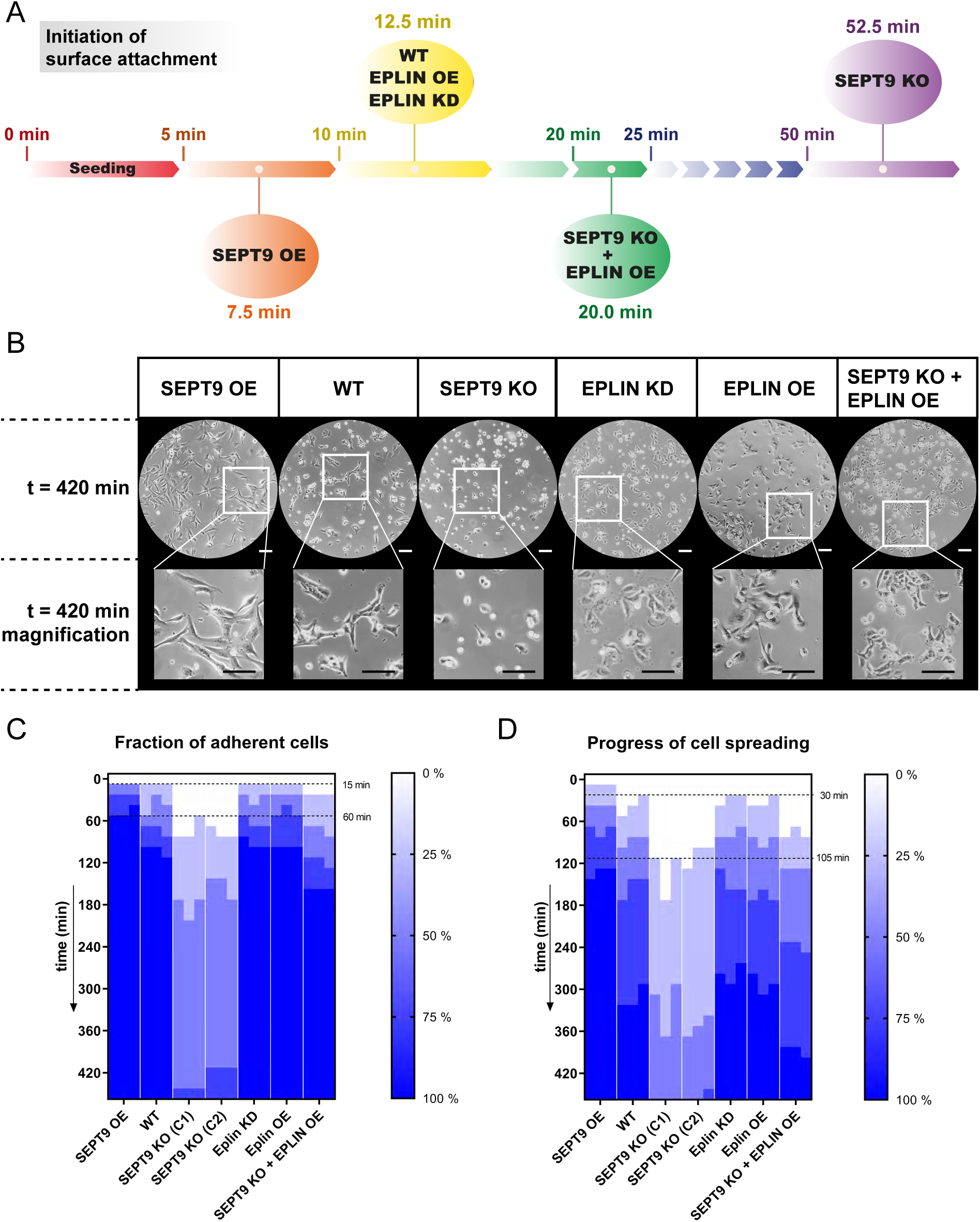
Reattachment and spreading are tightly regulated by SEPT9 and EPLIN. **(A)** The initiation of substrate reattachment was enhanced at high levels of SEPT9 and prolonged at low levels of SEPT9. Different expression levels of EPLIN did not influence this process. In absence of SEPT9, an OE of EPLIN could partially rescue the delayed attachment of SEPT9 KO cells. **(B)** The reattachment assay revealed a positive correlation of SEPT9 concentration with cell adhesion and spreading. EPLIN alone did not influence the attachment, however, could partially rescue the SEPT9 KO phenotype (10x objective, scale bar = 200 µm). **(C)** Heatmap representing the progress of cell-surface attachment. Based on light microscopy evaluation every 15-30 min, the reattachment was classified in five steps from 0 to 100 %. **(D)** Heatmap of cell spreading upon seeding. The assays in C and D were performed in triplicate. Cell adhesion and spreading was quantified by dividing the progress per replicate at each timepoint into five steps (0 %, 25 %, 50 %, 75 %, 100 %).

The mean time elapsed from seeding until substrate attachment of all cells was 105 min for WT, EPLIN OE and EPLIN KD cells. Elevated levels of SEPT9 shortened this process to 60 min, whereas a SEPT9 KO severely delayed the process to 12-16 h. These effects were even enhanced when the progress of cell spreading was additionally considered (Fig. 4D). To monitor spreading, the time was recorded until the normal morphology of at least 75 % of the cell culture was restored. This spreading time was significantly reduced to 120 min in SEPT9 OE compared to 300 min in WT cells. In SEPT9 KO cells, the spreading process was not properly measurable due to the extensive time delay (18-22 h) but both reattachment and cell spreading were accelerated by OE of EPLIN in these cells. Taken together, these results suggest an interplay between SEPT9 and EPLIN in the regulation of cell-surface interaction.

### SEPT9 influences the localization of EPLIN in cell protrusions

The interdependency of SEPT9 and EPLIN in cell adhesion and migration might indicate that SEPT9 influences the subcellular localization of EPLIN or vice versa. Encouraged by a recent study showing a pivotal role of EPLIN at the leading edge of migrating cells (Linklater et al., 2021), we focused on the subcellular localization of EPLIN and SEPT9 at cell protrusions of motile cells. Staining of endogenous EPLIN in stably GFP-SEPT9 expressing 1306 cells revealed a partial colocalization of EPLIN and SEPT9 at the tip of lamellipodia and along SEPT9 filaments (Fig. 5 A). Furthermore, EPLIN appeared in a dense belt-like sub-cortical network (Fig. 5 B, C). This EPLIN network around the cortex was restricted by SEPT9 towards the cell center. SEPT9 and EPLIN showed a partial overlap at this “restriction zone” where they formed a dense complex or filamentous structures. We analyzed this sub-cortical EPLIN structure in SEPT9 KO cells and WT fibroblasts and quantified the relative change in its thickness. SEPT9 KO cells had a 50 % reduced thickness of the EPLIN layer at the lamellipodium, whereas the diameter was enhanced 2.5-fold in SEPT9 OE cells (Fig. 5 D, E). In contrast, the SEPT9 distribution did not change upon EPLIN OE or KO (see below). EPLIN OE, however, induced the formation of eminently elongated filopodia along the whole cell membrane which we termed “paramecium phenotype” (Fig. 6A). The morphological changes ranged from spike-like membrane protrusions to a filament-like network formation within the cell. We quantified their length in cell lines displaying changing expression levels of SEPT9 and EPLIN (WT, EPLIN OE, EPLIN KD, SEPT9 KO and SEPT9 OE) (Fig. 6 B, C; see Suppl. Fig. S1D, E for the relative expression levels). Filopodia were visualized by actin staining. The average length of filopodia was increased 1.7-fold upon EPLIN OE compared to WT and a control cell line overexpressing GFP. The OE of SEPT9 had no significant influence on filopodia size. In contrast, the downregulation of SEPT9 or EPLIN equally decreased the length of filopodia. Overexpression of EPLIN in SEPT9 KO cells restored the length of the filipodia to wildtype levels. To test if the regulation of filopodia length depends on interaction of both proteins, we measured filipodia length upon OE of an GFP-EPLIN construct lacking the SEPT9-binding LIM domain. EPLIN_ΔLIM_ did not induce elongated filopodia. In contrast the mean size was reduced to levels found in in cells with a KD of EPLIN or a KO of SEPT9.

**Figure 5.**
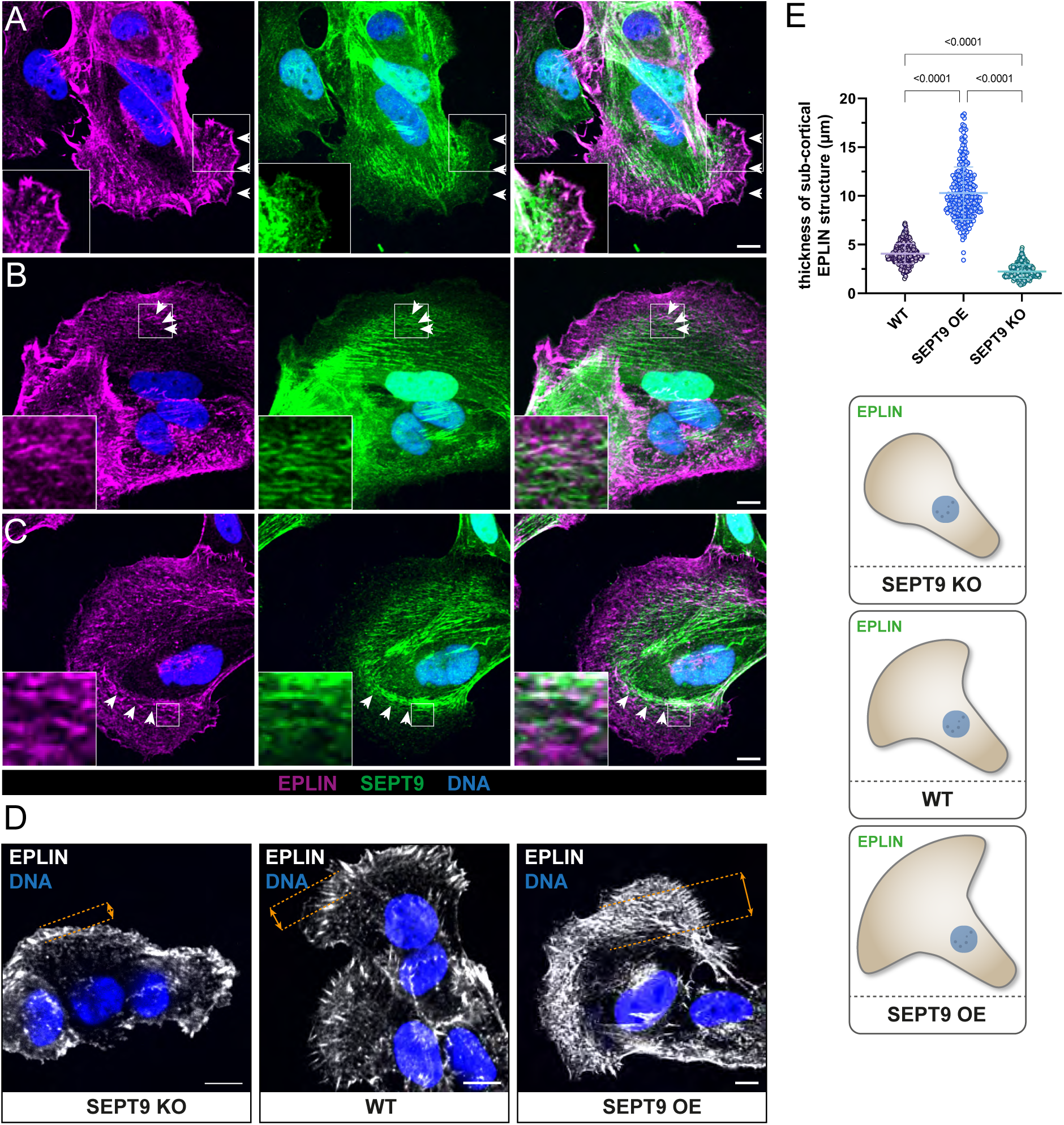
SEPT9 and EPLIN colocalize directly or are adjacent in cell protrusions. Fluorescence microscopy images show the localization of immunostained EPLIN in 1306 cells expressing GFP-SEPT9. **(A)** Cell protrusion where SEPT9 and EPLIN colocalized at the tip of a lamellipodium. EPLIN also localized along SEPT9 filaments in the cytoplasm. **(B, C)** Protrusions where the localization of EPLIN was restricted by the SEPT9 network. White arrows mark the location with a partial overlap of both proteins. **(D)** Cell protrusions showing the structural organization of EPLIN in relation to the SEPT9 expression level. All images in A-D represent average intensity projections from confocal microscopy (scale bar = 10 µm). **(E)** The quantification of the EPLIN thickness in cell protrusions (as indicated in D) shows a significant increase upon SEPT9 OE and a significant decrease upon SEPT9 KO. These findings are illustrated in the cartoon. Significance values were calculated by one-way ANOVA followed by Tukey’s multiple comparison test from three independent experiments, each with a sample size of n = 90 cells. The data are depicted as means ± SD.

**Figure 6.**
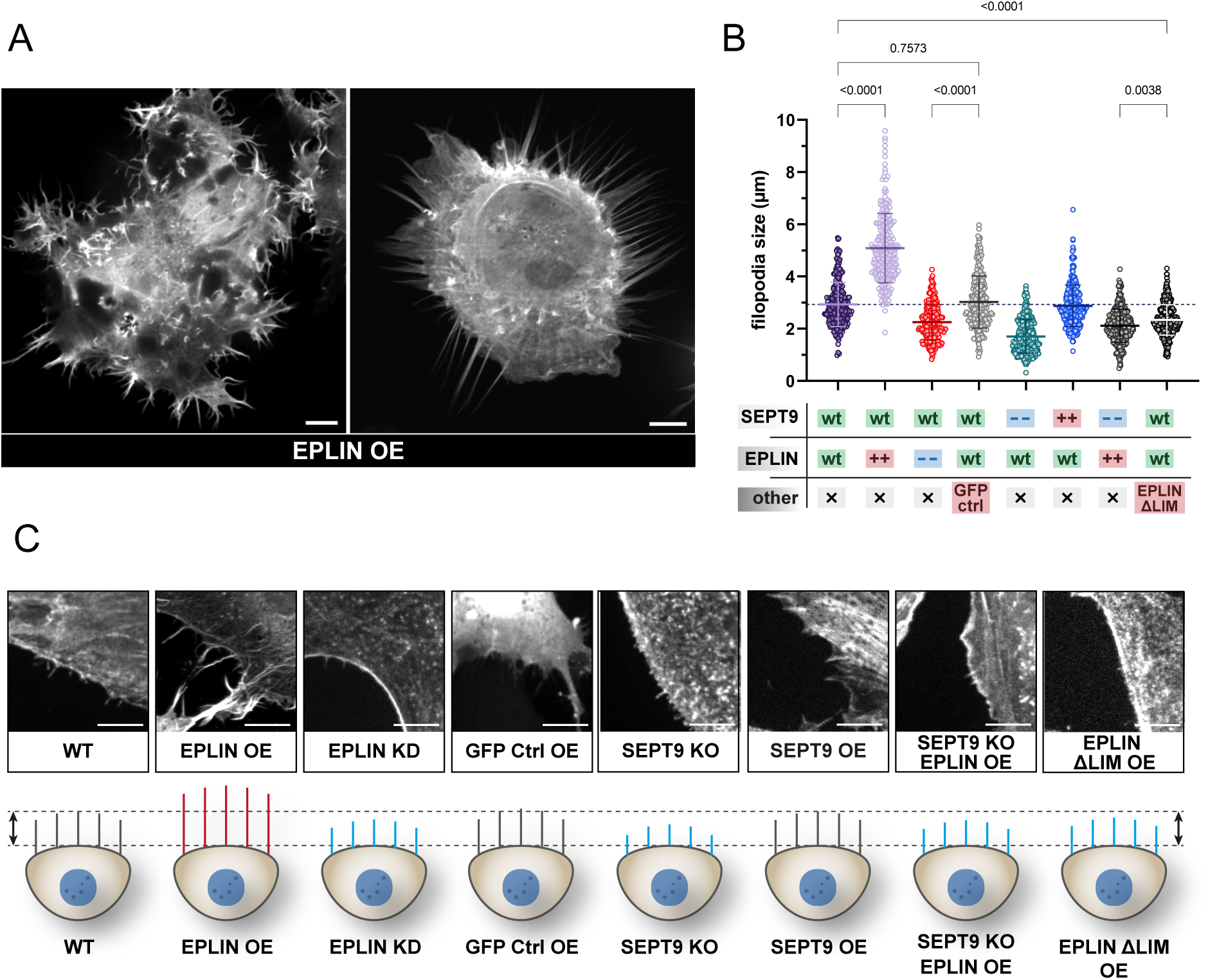
The size of filopodia is dependent on EPLIN and SEPT9. **(A)** The OE of GFP-EPLIN induced an enhanced formation of elongated filament-like or spike-like filopodia along the whole plasma membrane resulting in a paramecium-like appearance (scale bar = 10 µm). **(B)** Filopodia size of 1306 cells is dependent on EPLIN and SEPT9. An upregulation of EPLIN significantly increased filopodia size, whereas the individual downregulation of SEPT9 and EPLIN decreases their size. Neither the OE of EPLIN in SEPT9 KO cells nor the OE of an EPLIN mutant lacking the LIM domain showed elongated filopodia (“- -“, knockout or knockdown; “++”, overexpression). Analyses were performed 48 h upon transient overexpression of the respective construct. Significance values were calculated by one-way ANOVA followed by Šidák’s multiple comparison test from three independent experiments, each with a sample size of n = 80 cells. **(C)** Confocal microscopy images showing representative sections of filopodia stained by phalloidin (scale bar = 50 µm) and the corresponding schematic alteration. Filopodia in grey indicate a WT-like size, a decreased length is highlighted in blue and increased filopodia are presented in red (relative filopodia lengths are drawn to scale).

Changes in filopodia as part of the migration machinery raised the question, whether an impaired SEPT9-EPLIN interaction affects the overall migrative behavior of cells. Cell protrusions were tracked by actin staining by time lapse microscopy. Protrusions in WT cells showed a directed distribution at the leading edge (Video 2). Cells with EPLIN KD cells showed, in contrast, multiple, branched cell protrusions assembling simultaneously in multiple directions (Video 3). To confirm again that an impaired SEPT9-EPLIN interaction is responsible for this cellular defect, GFP-EPLIN_ΔLIM_, lacking the SEPT9 interaction site, was overexpressed and monitored in migrating cells. Time lapse microscopy showed the simultaneous presence of cell protrusions similar to those observed in EPLIN KD cells (Video 4). Taken together, these experiments confirmed that the interaction between EPLIN and SEPT9 is essential during migration to regulate the correct formation of protrusions at the cell tip.

### The interplay of SEPT9 and EPLIN regulates actin-based structures in fibroblasts

EPLIN was identified as an important member of the actin remodeling machinery (Collins et al., 2018; Taha et al., 2019). We next investigated the influence of SEPT9 and EPLIN on the actin cytoskeleton. In 1306 WT fibroblasts, the actin cytoskeleton consisted of transverse arcs or dorsal stress fibers at the cell cortex and ventral fibers towards the nucleus (Fig. 7, center panel). Upon OE of SEPT9 the number and thickness of ventral stress fibers that span the whole cell increased considerably. These bundled fibers were associated with the focal adhesion marker paxillin (Fig. 7 lower panel). In contrast, the KO of SEPT9 resulted in a complete loss of bundled actin filaments. The actin cytoskeleton in these cells consisted mainly of filamentous fibers along the plasma membrane (Fig. 7, upper panel). EPLIN did not affect stress fiber formation but influenced the distribution of actin filaments at the cortex. Overexpression of EPLIN translocated actin filaments towards the plasma membrane with an enhanced formation of filopodia. This effect was independent of SEPT9 (Fig. 7, right column). A reduction of EPLIN (Fig. 7, left column) led to a circular actin network with a reduced amount of ventral stress fibers that span the whole cell. A simultaneous upregulation of SEPT9 maintained the circular organization but again enhanced the bundling of paxillin associated actin filaments. The absence of both proteins abolished stress fibers and transverse arcs and left only thin filaments along the plasma membrane.

**Figure 7.**
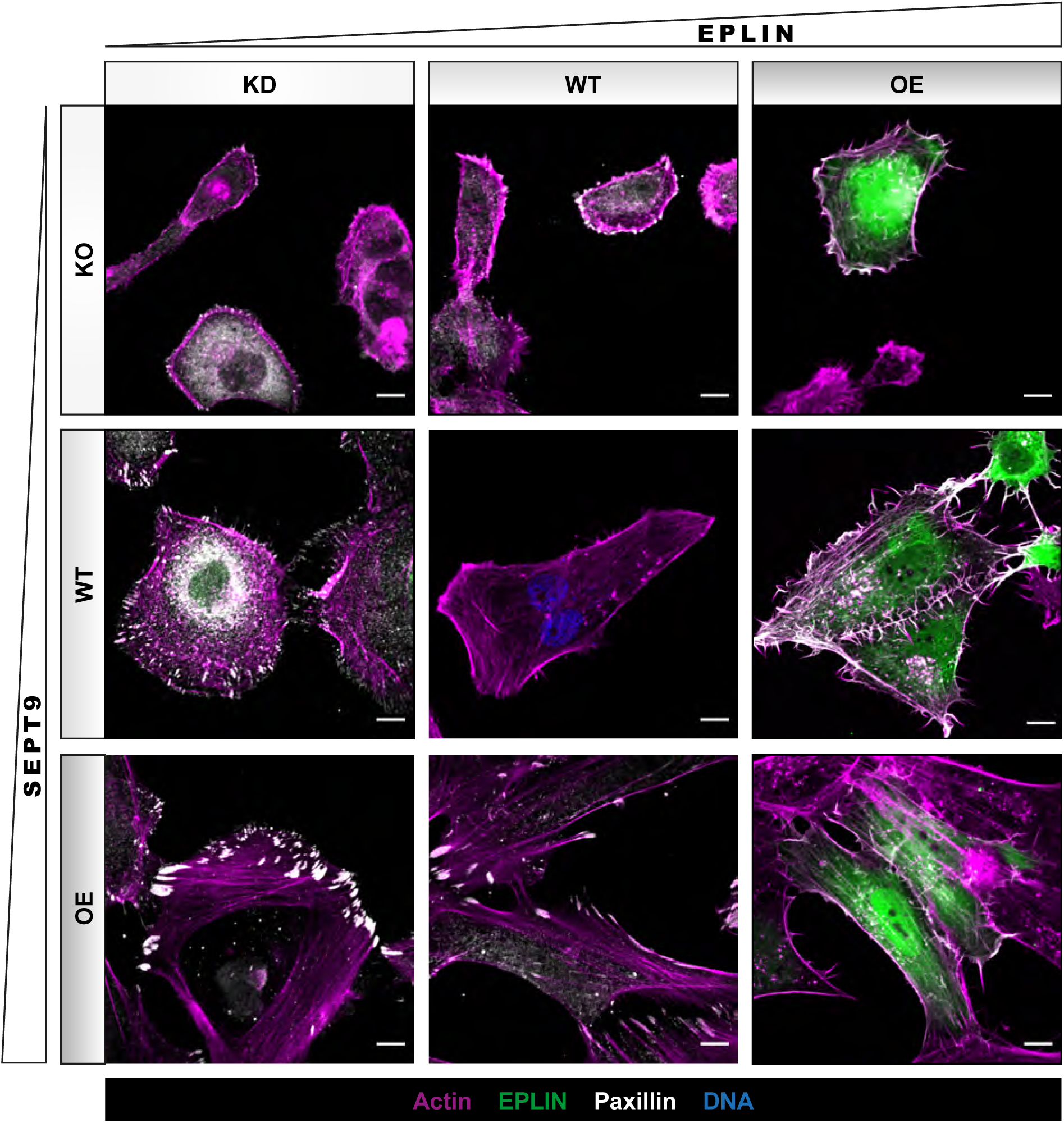
Actin organization in 1306 fibroblasts is strongly dependent on SEPT9 and EPLIN. In WT cells, actin (SiR-Actin) is organized in filaments and bundled fibers throughout the cells and reaches into focal adhesions. The KO of SEPT9 destabilized actin filaments and induced an accumulation along the plasma membrane. An additional KD of EPLIN slightly enhanced this effect, whereas the OE of GFP-EPLIN showed no severe impact on the actin structure. Compared to WT cells, the depletion of EPLIN induced a circular organization of actin filaments and rounding of cells. The simultaneous OE of GFP-SEPT9 stabilized actin filaments and induced circular bundles. The OE of EPLIN alone favored the formation of long filopodia with actin filaments that were enriched along the membrane, also upon additional OE of SEPT9. The OE of SEPT9 alone induced long, straight stress fibers that reached into large focal adhesions. Maximum projections of confocal microscopy images are presented (scale bar = 10 µm).

Taha *et al*. suggested that EPLIN interacts with the Arp2/3 complex to regulate protrusion dynamics (Taha et al., 2019). Furthermore, an influence of Arp2/3 on cell adhesion and protrusion formation was reported (Beckham et al., 2014). However, we could not detect any obvious effects of the Arp2/3 inhibitor CK666 on the EPLIN- or SEPT9-induced alterations of actin architecture or protrusion formation (Suppl. Fig. S4). The mobility of CK666 treated 1306 WT decreased by 76 %, resulting in nearly immobile cells as detected by a Boyden chamber assay (Suppl. Fig.S5). However, SEPT9 OE and EPLIN KD cells were less sensitive to CK666 treatment. Compared to DMSO-treated cells, the migration rate of both cell lines decreased by less than 10 %. EPLIN partially localizes along septin and actin filaments but is also found in the tip of lamellipodia and together with paxillin in focal adhesions. The downregulation of EPLIN in 1306 fibroblasts reduced not only the size of focal adhesions but also translocated paxillin-rich structures from the membrane towards the cytoplasm (Fig. 8A, left column). In contrast, the upregulation of EPLIN strengthened paxillin structures at the plasma membrane (Fig. 8A, right column). Both up- and downregulation of SEPT9 led to a preferential localization of paxillin at the plasma membrane with smaller adhesions upon SEPT9 KO and larger adhesions upon SEPT9 OE (Fig. 8A, right column). Using paxillin as marker, we subsequently quantified the size of focal adhesions upon up- or downregulation of EPLIN and SEPT9 (Fig. 8 B, C, D). The OE of SEPT9 increased the size of focal adhesions of 4.9 µm compared to the WT (3.7 µm). In contrast, the KO of SEPT9 led to a reduction in focal adhesion length (2.1 µm). In accordance with the increase and decrease of SEPT9, the KD of EPLIN reduced (2.5 µm) and its OE elongated (4.2 µm) the size of adhesions, however to a lesser extent than SEPT9.

**Figure 8.**
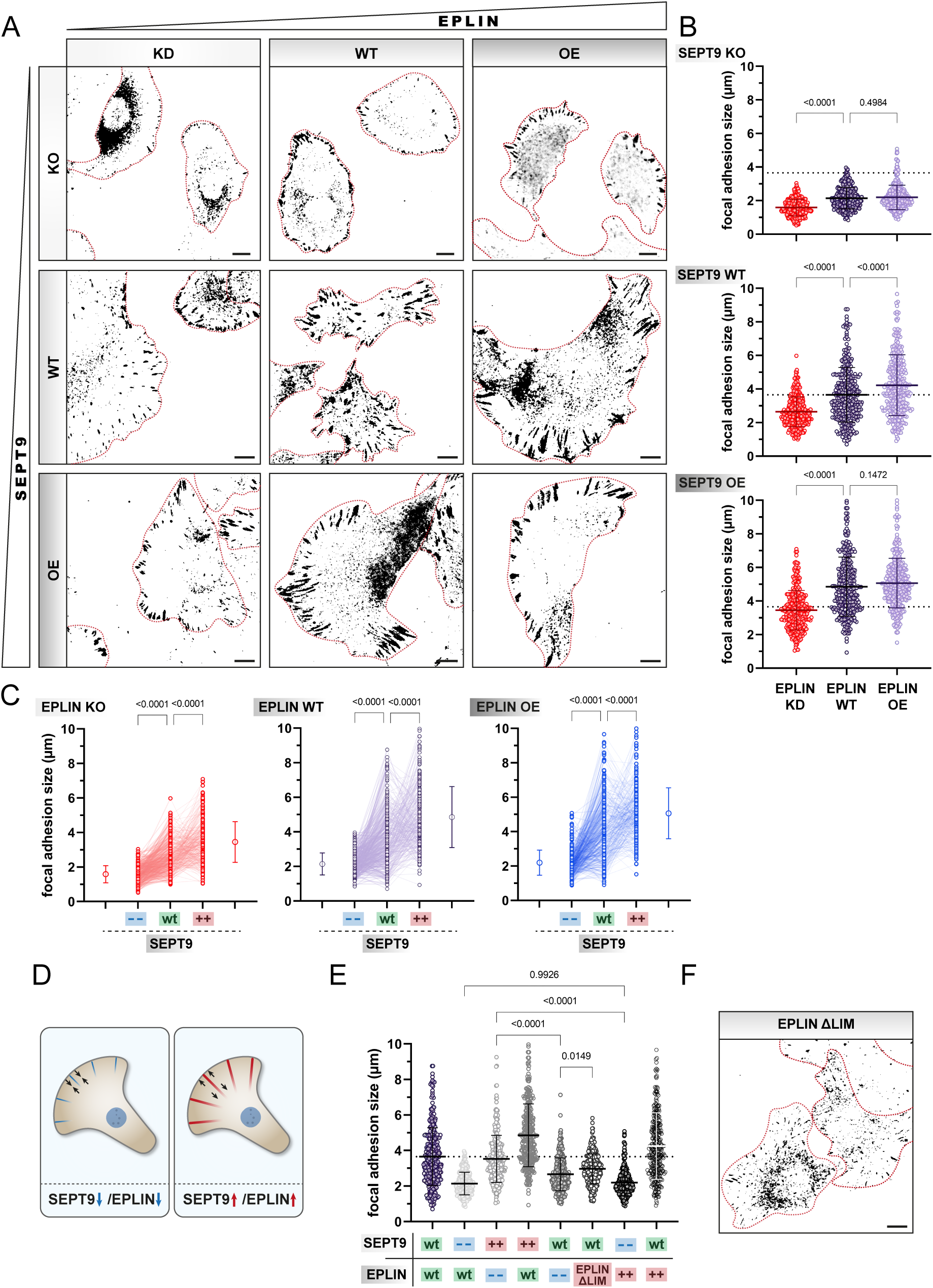
The size and shape of focal adhesions is dependent on SEPT9 and EPLIN. **(A)** Immunostaining of paxillin upon up- and downregulation of EPLIN and SEPT9. For better visibility paxillin is shown as inverted image in black. Individual cells are outlined in red (scale bar = 10 µm). Graphical representation of focal adhesion length in dependence of SEPT9 **(B)** and EPLIN **(C)** (“- -“, knockout; “+ +”, overexpression). Significance values were calculated by one-way ANOVA followed by Šidák’s multiple comparison test from three independent experiments, each with a sample size of n = 100 cells. **(D)** Scheme showing the reduction and increase of focal adhesions in dependence of SEPT9 or EPLIN. **(E)** Quantification of the EPLIN_ΔLIM_ mutant revealed reduced focal adhesion size, In contrast, full length EPLIN in WT cells increased the size of focal adhesions. (“- -“, knock down; “+ +”/red field, overexpression). Significance values were calculated by one-way ANOVA followed by Dunnett’s multiple comparison test from three independent experiments, each with a sample size of n = 100 cells. Quantitative data are depicted as means ± SD. **(F)** Immunostaining of paxillin upon overexpression of EPLIN_ΔLIM_. For better visibility paxillin is shown as inverted image in black. Individual cells are outlined in red (scale bar = 10 µm).

To identify potential synergistic effects of EPLIN and SEPT9, we additionally measured the size change in a SEPT9 KO/EPLIN KD as well as in a SEPT9 OE/EPLIN OE cell line (Fig. 8 B). The double depletion reduced the size of focal adhesions further (1.6 µm) whereas the double OE had no severe impact (5.1 µm) compared to the single protein OEs. Only the OE of SEPT9 in EPLIN KD cells (3.5) could restore a focal adhesion length comparable to WT. The OE of EPLIN in SEPT9 KO cells (2.2 µm) was not sufficient to induce larger adhesions. Their size remained at the same length as in SEPT9 KO cells with WT levels of EPLIN.

Migration defects were observed in cells with impaired SEPT9-EPLIN interaction including SEPT9 KO cells and cells expressing the EPLIN_ΔLIM_ construct. In comparison to full length EPLIN with increased paxillin structures, EPLIN_ΔLIM_ had an opposite effect by decreasing the size of focal adhesions (3.0 m) (Fig. 8E). Although this mutant of EPLIN could not reduce the focal adhesion size to the same extent as the complete KO of SEPT9 or KD of EPLIN, a significant decrease was measured compared to the WT (Fig. 8F).

Taken together, by identifying a correlation between length variation in paxillin structures and impaired migrative behavior of 1306 cells we were able to show that the SEPT9-EPLIN interaction is essential in the regulation of migration via focal adhesions.

## Discussion

### SEPT9 and EPLIN act in concert to regulate cell migration and surface attachment

We previously identified EPLIN in a proteomics screen as a novel interaction partner of SEPT9 (Hecht et al., 2019). We confirm here that the G-domain of SEPT9 is sufficient to mediate the zinc-dependent binding to the LIM domain of EPLIN *in vitro*. We show further that the EPLIN localization throughout the whole cell cycle is strongly dependent on SEPT9.

The positive correlation between cell movement and the expression level of SEPT9 suggested a crosstalk between the septin cytoskeleton and proteins that actively generate forces to induce changes in cell shape and motility. Our nearly immobile SEPT9 KO cells emphasize the importance of a balanced SEPT9 regulation for cell migration. The idea of a SEPT9-dependent mechanosensitive regulation is supported by the work of Yeh *et al*., who showed that SEPT9 is upregulated in response to soft substrates, whereas stiff matrices induced a downregulation of SEPT9 in endothelial cells (Yeh et al., 2012). Instead of varying the substrate stiffness and analyzing endogenous SEPT9 levels, we varied the level of SEPT9 and studied adhesion and spreading of these cells. The time between cell seeding and the initiation of surface attachment was reduced (- 40 %) by high SEPT9 levels and severely prolonged (+ 320 %) upon SEPT9 KO. These results were accompanied by a 3-fold faster attachment and spreading process in SEPT9 OE and a more than 3-fold slower rate in SEPT9 KO cells. We thus propose a direct mechanistic link between the adhesive capability of a cell and the level of SEPT9, and suggest that that EPLIN as SEPT9 interaction partner represents this link. The OE or KD of EPLIN did not significantly alter the surface attachment compared to WT cells. The severe delay of cell-surface interaction in SEPT9 KO cells could partially be rescued by elevated levels of EPLIN. The deletion of EPLIN is associated with a destabilization of the cadherin complex (Abe and Takeichi, 2008; Taguchi et al., 2011). Elevated levels of EPLIN were shown to promote the formation of linear actin filaments by inhibiting the Arp2/3-mediated branching (Maul et al., 2003). This explains why the OE of EPLIN enlarges the cell surface area of SEPT9 KO cells. Providing a larger contact area between cell and substrate in combination with an enhanced filament formation rate reduces the time required for cell attachment and spreading. However, our data suggest that initiation and progression of cell reattachment are not exclusively regulated by EPLIN.

Cells treated with the Arp2/3 inhibitor CK666 showed no significant alteration in actin filaments upon up- or downregulation of EPLIN and SEPT9, respectively. In accordance with these observations, treatment with CK666 had only a minor effect on the motility of the cells. Our data thus contradict the model that EPLIN acts on actin filaments exclusively through the regulation of Arp2/3. Instead, we suggest that the MAPK/ERK (mitogen-activated protein kinases/extracellular signal-regulated kinases) pathway is involved in EPLIN/SEPT9-regulated cell migration. Our MS screen identified two components of this pathway, MLCK and MYPT1, as interaction partners of SEPT9 (Hecht et al., 2019). Both proteins participate in the formation and maintenance of lamellipodia and actin stress fibers (Ghosh et al., 2021; Joo and Yamada, 2014). This observation is supported by the finding that a downregulation of SEPT9 is associated with a reduction of activation of the MEK/ERK pathway in Glioblastoma cells (Xu et al., 2018). EPLIN, in turn, is a direct downstream phosphorylation target of ERK (Han et al., 2007).

We propose that the regulation of cell adhesion and migration through EPLIN and SEPT9 is based on three major features: a) remodeling rate of the actin cytoskeleton, b) bundling and stabilization of actin filaments and c) sensing of cellular tension. EPLIN was shown to act as a mechanosensor at adherens junctions (Taguchi et al., 2011). In our model, mechano-sensitivity of EPLIN would rely on the presence of SEPT9 as a modulator of cellular tension and as spatial guide for EPLIN. Through the direct interaction of EPLIN with the cadherin-catenin complex, intracellular forces are connected to the adhesive machinery (Chervin-Pétinot et al., 2012).

Our model for the following scenarios (SEPT9 KO, WT, SEPT9 OE) may explain how a balanced level of EPLIN regulates the remodeling of the actin cytoskeleton and thereby overall cellular dynamics. At WT SEPT9 levels, actin filaments are organized in a dynamic process providing stability and flexibility to the cell. In presence of EPLIN, its mechanosensory function balances the adhesion machinery and the Arp2/3 mediated actin-branching (Fig. 9A, middle). Upon EPLIN KD the cell is unable to sense the level of extracellular forces or cytoskeletal tension, which enhances actin branching at the cell front and thereby cell migration. The deletion of SEPT9 results in a reduction of bundled actin filaments and of stress fibers. The reduced cellular tension may be sensed by EPLIN to limit the dynamics of the actin cytoskeleton (Fig. 9A, left), resulting in a low motility rate (Fig. 3C). Consequently, the KD of EPLIN and the resulting loss of its mechanosensing capability in SEPT9 KO cells induces an enhanced motility (Fig. 3C) even in absence of stress fibers and actin filament bundles. SEPT9 OE shifts the equilibrium of actin towards bundled filaments with high tension. The mechanosensing of a high-tension state through EPLIN promotes cell-surface adhesions to facilitate migration (Fig. 9A, right). In SEPT9 OE cells, the KD of EPLIN does not induce an increase in cell motility (Fig. 3C). The stabilization and mechanical support of actin bundles by SEPT9 limits actin branching and thereby inhibits excessive cell migration as observed upon EPLIN KD in WT or SEPT9 KO cells.

**Figure 9.**
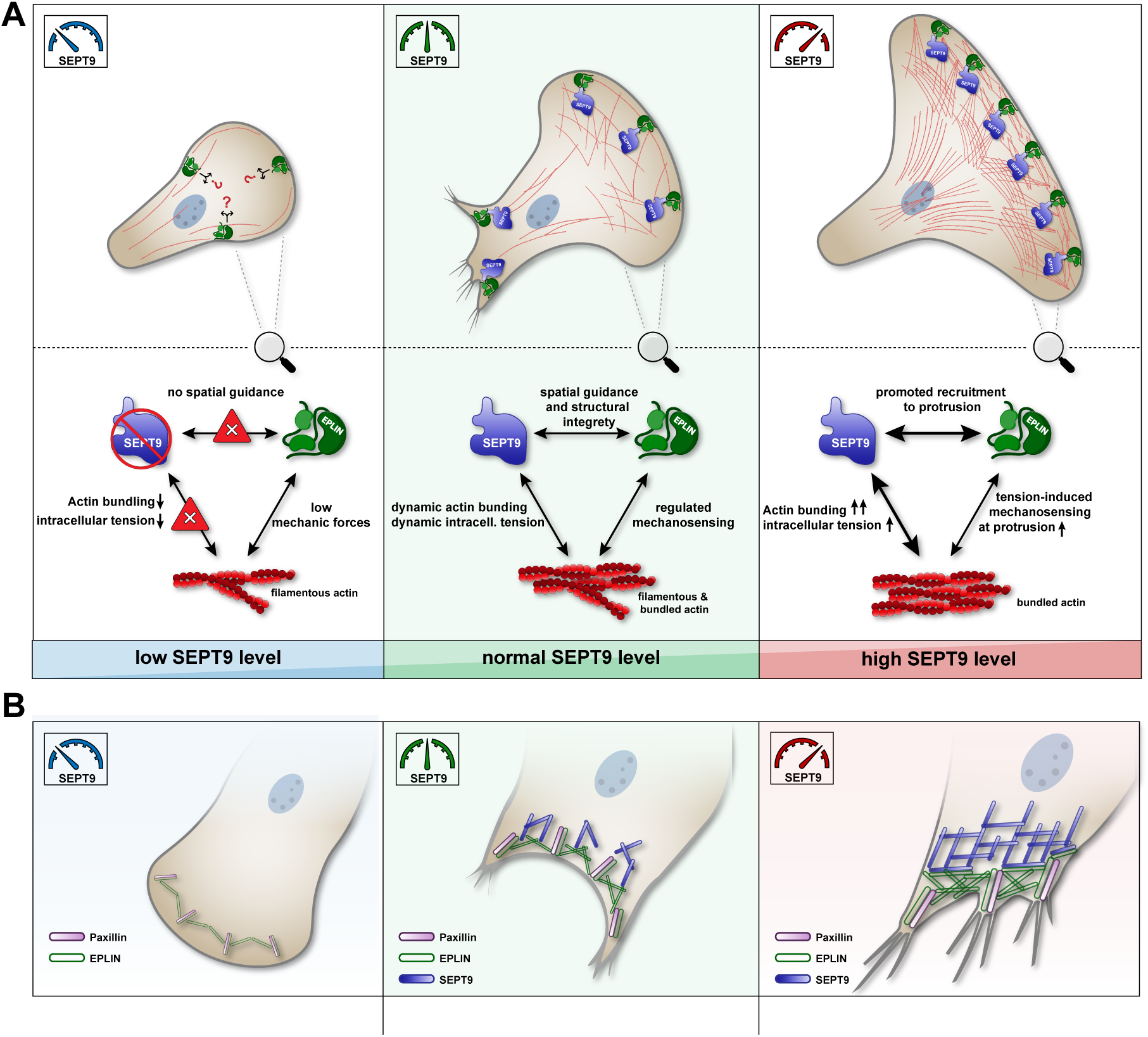
Model of the mutual SEPT9 and EPLIN interplay in cell migration and focal adhesion regulation. **(A)** In WT cells, SEPT9 defines the mechanical properties of actin through its bundling activity. Regions of high tension (e.g., at the tip of migrating cells) are sensed by EPLIN to recruit the adhesion complex. SEPT9 may act as scaffold for the localization of EPLIN along the lamellum or recruits EPLIN-binding proteins to the front. Without SEPT9 (left panel), actin is less bundled and predominantly present in its filamentous state. Low tension actin filaments cannot recruit EPLIN correctly to the leading edge. The enrichment of EPLIN at the cell tip is reduced, as absent SEPT9 cannot trigger its spatial recruitment. Elevated levels of SEPT9 (right panel) promote the bundling of actin (high tension state) and the recruitment of EPLIN towards the cell front. Increased mechanical forces in the cell are sensed by EPLIN and “hyperactivate” the adhesion machinery. **(B)** The core multi-protein focal adhesion complex is assembled independently of SEPT9. The presence of SEPT9 is, however, responsible for the stabilization and growth of this complex mediated through actin and its direct interaction with EPLIN. Elevated level of SEPT9 (right panel) enhance the formation of octameric septin building blocks, which promote actin bundling and localize EPLIN to the sites of focal complexes. EPLIN presumably induces the recruitment of additional components such as paxillin, thereby promoting larger FAs. Reduced levels of SEPT9 (left panel) limit the amount of EPLIN at FAs, prevent the addition of further adhesion proteins. The complex is therefore limited to the minimal “core components”, which allow the formation only of weak FAs.

### SEPT9 modulates the organization and localization of focal adhesions through EPLIN

At the lamellum, where densely structured SEPT9 is separated from EPLIN, we observed a partial overlap along individual filaments. Cells with an elevated level of SEPT9 showed a denser network of EPLIN at the cell tip, whereas the absence of SEPT9 reduced the amount of EPLIN at the cell front and restricted cell migration. We propose that SEPT9 does not only act as a mediator of actin filament bundling but also promotes the accumulation of EPLIN at the leading edge. Colocalization studies in SEPT9 OE cells suggested, that the SEPT9-rich lamellum directly binds to EPLIN and and regulates its local concentration. In epithelial cells, EPLIN responds to mechanical forces and is associated to the *zonula adherens* through its interaction with α-catenin (Taguchi et al., 2011). Previous studies connected the intracellular expression level of SEPT9 with the morphology of focal adhesions (Fuechtbauer et al., 2011; Dolat et al., 2014). By showing an increase in focal adhesion number and length with increasing levels of SEPT9 and EPLIN our data incorporate EPLIN in this pathway. In absence of SEPT9 or in absence of EPLIN, focal adhesions were significantly reduced in size. An additive reduction in focal adhesion size was measured in double depleted cells (EPLIN KD in SEPT9 KO cells). This suggests that the presence of both proteins is required to maintain a stable focal adhesion complex.

Similar to cells with EPLIN KD, cells overexpressing EPLIN_ΔLIM_, lacking the SEPT9 interaction site, showed on the one hand a reduced focal adhesion size and on the other hand a re-localization from the membrane towards the cytoplasm. Taken these findings together, we propose that EPLIN binds to the focal adhesion machinery via its known interaction partners α-catenin (Chervin-Pétinot et al., 2012) and paxillin (Kasai et al., 2018) independently of SEPT9. However, the local concentration of EPLIN seems to depend on the intracellular amount of SEPT9. With increasing levels of SEPT9, the septin cytoskeleton at the lamellum recruits actin filaments, promoting the local concentration of EPLIN. This enhanced local enrichment of EPLIN at the leading edge would then initiate a cascade to recruit additional binding partners of the adhesome complex (Fig. 9B, right). A reduction of endogenous SEPT9 restricts the formation of actin bundles and thus limits the localization of EPLIN to focal adhesions.

Lowered concentrations of EPLIN cannot recruit additional focal adhesion components, resulting in smaller and fewer adhesion complexes. Shrunk adhesive structures provide less contact area to the surface and thereby weaken the adhesive and migrative behavior of SEPT9 KO cells (Fig. 9B, left).

Motility and adhesion are the outcome of the collective behavior of hundreds of different components. The list of the participating proteins is most likely still not complete and their roles are far from understood. Our data show how SEPT9 through its interaction with EPLIN and actin contributes to both processes.

## Materials and Methods

### Plasmids, cloning and siRNA

All used and generated plasmids are summarized in Table 1. Cloning was performed using standard procedures. The ORF of EPLIN isoform alpha was PCR amplified from Addgene plasmid #40928 to generate plasmid #1. All other EPLIN constructs were derived from this plasmid. All SEPT9 constructs were derived from plasmid #4 (kind gift from E. Spiliotis, Drexel University, PA, USA). The iRFP ORF was PCR amplified from Addgene plasmid #45457.

**Table 1.** List of plasmids. pAC-GST and pES are an in-house constructed, pET15A based plasmids for the expression of His_6_ tagged proteins in E. coli.

|  | Construct | Vector backbone | Insert | Source |
| --- | --- | --- | --- | --- |
| #1 | EGFP-EPLIN | pLenti-CMV-GFP-Puro | fl (isoform alpha) | This study |
| #2 | EGFP-EPLIN LIM | pLenti-CMV-GFP-Puro | aa 228-288 | This study |
| #3 | EGFP-EPLIN (ΔLIM) | pLenti-CMV-GFP-Puro | Δaa 228-288 | This study |
| #4 | mRuby-EPLIN | mRuby2-C1 (Addgene #55911) | fl | This study |
| #5 | EGFP-SEPT9 | pEGFP-C2 | fl | *E. Spiliotis* |
| #6 | GST-EPLIN | pGEX2T | fl | This study |
| #7 | GST-EPLIN LIM | pGEX2T | aa 228-288 | This study |
| #8 | GST-SETP9 | pAC-GST | fl | This study |
| #9 | His-EPLIN | pES | fl | This study |
| #10 | His-SEPT9 | pES | fl | This study |
| #11 | His-SEPT9 ΔC | pES | aa 1-567 | This study |
| #12 | His-SEPT9 ΔN | pES | aa 295-586 | This study |
| #13 | His-SEPT9 ΔN ΔC (G domain) | pES | aa 295-567 | This study |
| #14 | pSpCas9(BB)-2A-iRFP670 | pSpCas9(BB)2A-GFP (Addgene #48138) | riRFP670 | This study |
| #15 | pSpCas9(BB)-2A-GFP-Ex4B | pSpCas9(BB)2A-GFP (Addgene #48138) | SEPT9 Exon4 sgRNA | This study |
| #16 | pSpCas9(BB)-2A-iRFP-Ex6E | #14 | SEPT9 Exon 6 sgRNA | This study |

All used PCR primers are listed in Suppl. Table 1. siRNA targeting the LIM domain of EPLIN (# SASI_Hs02_00326071; TATTGTAAGCCTCACTTCAA) and a scrambled control siRNA (#SIC001) were obtained from Sigma-Aldrich.

### Expression of recombinant proteins and in vitro pulldown experiments

All recombinant proteins were expressed in *E. coli* BL21DE3 in super broth (SB) medium. Protein expression was induced with 0.1 mM IPTG (supplemented with 2 % v/v Ethanol for SEPT9 expression constructs) at an O.D._600_ of 0.8 and protein expression was conducted for 5 h (EPLIN constructs) or overnight (SEPT9 constructs) at 18 °C. The cells were harvested by centrifugation and stored at -80 °C until further use. Cell lysis was performed in appropriate buffer (see below) by treatment with 1 mg/ml lysozyme and ultrasound and cell debris was removed by centrifugation (40.000 xg, 10 min). For His_6_-tagged proteins, IMAC buffer A (50 mM KH_2_PO_4_, 20 mM Imidazole, 300 mM NaCl, pH 7.5) supplemented with e-complete protease inhibitor cocktail (Roche) was used as lysis buffer. These proteins were purified using a 5 ml HisTrap Excel column (Cytiva) mounted on an Äkta Pure chromatography device (Cytiva). Elution was carried out with a step gradient consisting of consecutive steps of 15%, 30% and 100% IMAC elution buffer (50 mM KH_2_PO_4_, 200 mM Imidazole, 300 mM NaCl, pH7.5). The proteins were subsequently buffered in PBS using a PD10 desalting column (Cytiva), concentrated and used for pulldown. GST-fusion proteins were directly lysed in PBS supplemented with e-complete protease inhibitor cocktail as described above. The resulting extract was immediately used for pulldown experiments. For each sample, 50 µl of PBS equilibrated Glutathione Sepharose beads (Cytiva) were incubated for 20 min with 500-1000 µl extract of the respective GST-tagged protein (or 500 µl of 2 µM purified GST for controls) under rotating agitation.

After washing with PBS, nonspecific binding sites were blocked by incubation with 1.5 µM BSA for 15 min. Purified His_6_-tagged proteins were added at a concentration of 1.5 µM and incubated for 30 min. Unbound proteins were washed away with PBS prior to an elution step with GST-elution buffer (50 mM Tris pH 8.0, 10 mM reduced Glutathione) for 10 min. Samples of the elutates and input proteins were subsequently analyzed with SDS-PAGE and Western blot, respectively. SDS-PAGE was performed using Bolt 4-12 % BisTris gradient Gels (Invitrogen) according to the manufacturer’s instructions. For Western blot, proteins were transferred onto nitrocellulose membrane which was subsequently blocked by 3 % skimmed milk. All employed antibodies and their applied dilutions are summarized in Suppl. Table 2.

### Cell culture and immunofluorescence

The human dermal fibroblast cell line 1306 was used for all experiments. Cells were routinely cultured at 37°C and 5 % CO_2_ in a humidified incubator in DMEM supplemented with 10 % FBS (both from Gibco). Antibiotics were only added for selection or maintenance of stable cell lines.

For transfection, the cells were seeded the day before to reach a confluency of 70-80 % on the day of transfection and the medium was renewed 30 min prior to transfection. Transfection of plasmid DNA was performed with Lipofectamine 3000 (Invitrogen / Life Technologies) according to the manufacturer’s instructions except that cells were transfected for 6 h before fresh medium was added. Typically, the cells were allowed to recover and express proteins for 48 h. Stable cell lines were selected by antibiotic treatment for two weeks with 1 µg/ml Puromycin (Formedium) or 750 µg/ml Geneticin G418 (Formedium). The antibiotic concentration was then lowered to 0.25 µg/ml or 250 µg/ml, respectively, and the cells were further cultivated under these conditions. For a mix-clone cell line, fibroblasts were further expanded, and the was modification was verified via Western blotting or immunofluorescence. To generate a monoclonal cell line, serial dilution was applied to singulate antibiotically selected cells in 96-well plates. The regrowth of individual clones was monitored until the appearance of larger colonies, and the dilution procedure repeated if necessary.

Duplex siRNA was transfected using Lipofectamine RNAiMAX (Invitrogen) according to the manufacturer’s instructions using 30-40 pmol siRNA per well of a 6-well cell culture plate. For protein knockdown, the Lipofectamine-siRNA mix was incubated for 6 h on the cells prior to media change and the cells were then incubated additional 48 h prior to experimental use. For co-transfection of siRNA and plasmid DNA, the RNA was already removed 4 h post-transfection. The cells were then allowed to regenerate for 2 h prior to a second transection of DNA with Lipofectamine 3000 overnight. The growth medium was renewed the day after, and cells incubated for further 36-48 h.

For live-cell microcopy, cells were seeded at least 24 h before microscopy in an 8-well culture chamber with a cover slip bottom (Sarstedt). 4-12 h prior to microscopy, the growth medium was removed and replaced by FluoroBrite DMEM (Invitrogen) supplemented with 10 % (w/v) FBS and 1 % (v/v) GlutaMAX (Invitrogen). If applicable, dyes to stain DNA or actin were added according to the manufacturer’s recommendations. SpyDNA (Spirochome) was used at a dilution of 1:4000 and SiR-Actin (Spriochome) at 1:5000-1:10,000 with an incubation time of 15 h prior to microscopy.

Immunofluorescence was performed either on sterile coverslips placed in 6-well plates or in 8-well cell culture chambers with cover slide bottom (Sarstedt). The day before, 3.0 x 10^5^ or 2.5 x 10^4^ cells were seeded per 6-well or 8-well, respectively. Cells were washed once with pre-warmed PBS immediately followed by covering the surface with 4 % (w/v) PFA for 10 min. Fixed cells were washed with PBS and permeabilized with 0.1 % TritonX100 in PBS for 20 min. Blocking was performed with 5 % (w/v) BSA in PBS for 45 min. Primary antibodies were diluted in Antibody Diluent (Dako) and a droplet of 25 µl or 50 µl applied on the sample for an 8-well or a coverslip upside down, respectively. The staining was performed for 1-4 h at RT or overnight at 4 °C in a humidified environment. Co-staining with multiple antibodies was performed in a single step, by mixing up to three antibodies in Antibody Diluent. Unbound antibodies were removed by washing with PBS before secondary antibodies were applied accordingly for 1 h at RT in dark environment (typical dilution 1:500 in Antibody Diluent). A list of all employed antibodies is provided in Suppl. Table 2. After washing, cells were coated with mounting medium (Dako), covered with a cover slip (8-well slide chamber) or a glass slide (cover slips) and stored at least for 6 h at 4 °C in the dark until microscopy.

### Generation of a SEPT9 CRISPR/Cas9 knock out cell line

We utilized the error-prone nature of NHEJ (non-homologous end joining) leading to frameshifts in the coding sequence to generate a SEPT9 knock out cell line in 1306 fibroblasts following an already established protocol (Ran et al., 2013) with some modifications. To identify specific and efficient target sites, the CRISPR design tool in Benchling (https://www.benchling.com/) was used for identification and optimization of small guide RNAs (sgRNAs/ gRNAs). The predicted efficiency was based on a scoring system that calculated the on-target and off-target value of each gRNA (Doench et al., 2016). An off-target value above 65, and an on-target above 60 were considered as good guides. To enhance the cleaving efficiency, single gRNAs were individually guiding Cas9 to the beginning of Exon 4B and to the end of Exon 6E of SEPT9, respectively (template sequence: ENSG00000184640). For cloning into the pSpCas9(BB)2A-GFP plasmid (Ran et al., 2013), *BbsI* overhangs were added to the respective sequences. The resulting sgRNA sequences are shown in Suppl. Table 3.

We replaced the ORF of the GFP marker in the pSpCas9(BB)2A-GFP plasmid (Addgene plasmid #48138) by the ORF of iRFP670, resulting in pSpCas9(BB)2A-iRFP670. The forward and reverse sgRNA oligonucleotides were annealed and the resulting sgRNA insert targeting Exon 4B was cloned into *Bbs1* restricted pSpCas9(BB)2A-GFP and the sgRNA insert targeting Exon 6E was cloned into *Bbs1* restricted pSpCas9(BB)2A-iRFP670. Annealing and cloning were performed as described elsewhere (Ran et al., 2013).

The plasmids were transfected into 1306 fibroblasts and sorted by FACS 48h post transfection. GFP and iRFP670-positive cells were singulated and sorted into 96-well plates previously seeded with Mitomycin C-treated MEF feeder cells (kindly provided by Prof. M. Füchtbauer, Aarhus, Denmark). Upon colony formation, the individual clones were expanded and the DNA was isolated using the QuickExtract DNA Extraction Solution (Lucigen). A successful homozygous genomic modification was confirmed by PCR (data not shown) and the absence of respective translation product by Western blotting (Suppl. Fig. S2). We used herein two selected clones with identical properties (Clone C1 (2c13) Clone 2 (6c29)).

### Transwell migration assay (Boyden chamber assay)

Non-confluent cells were detached from a culture dish and counted. 50,000 cells were resuspended in 100 µl prewarmed DMEM without FBS and transferred into a 8 µM PET tissue culture plate insert (BrandTech). The insert was placed into a well of a 24 well cell culture plate containing 700 µl prewarmed DMEM supplemented with 10 % FBS. The cells were allowed to attach and migrate for 20-24 h at 37 °C. The following day, the insert was washed twice with PBS inside and outside prior to fixation with 4 % (w/v) PFA in PBS for 20 min. After two additional washing steps with PBS, the cell nuclei were stained with a 1:2000 dilution of SPY-DNA dye (Spirochrome) in antibody diluent (Dako). SPY-DNA 505, 555 or 650 were used depending on the fusion protein in the respective cell line.

The documentation of migrated and non-migrated cells was performed in two consecutive steps by fluorescence microscopy. First, the transwell inserts were transferred to a glass slide and at least ten images at random positions were taken. Non-migrated cells on the top layer of the insert membrane were then removed by a cotton swab and ten additional images were taken. The function “Find Maxima” in FIJI was used to automatically identify nuclei in an image. The threshold to exclude background signals was set to a prominence > 5,000-20,000 ensuring that only one datapoint per nucleus was generated. The ratio of migrated cells to the total cell count was used to calculate the migratory behavior of the respective cell line.

### Surface reattachment and cell spreading assays

Cell reattachment and spreading were evaluated simultaneously from the same dataset. Non-confluent cells were detached with trypsin, counted, and seeded to a density of 2.0-3.0 x 10^5^ cells in a 6-well plate. Immediately, a photo (t = 0 min) of a random position was taken through the 10x objective on a routine light microscope (Zeiss Axiovert 40C) using a OnePlus 8T camera at 12 MP (1,6 µm/px) and the plate was placed back in a CO_2_-Incubator at 37°C. Cell morphology was documented accordingly in triplicate every 15-30 min for a total of 450 min. To evaluate the adhesion of cells, the culture plate was gently rocked back and forth under the microscope.

The level of adhesion was classified in five steps from “none” to “complete” (0 %, 25 %, 50 %, 75 %, 100 %), defined by the fraction of cells adhering to the plastic surface. The progress of spreading was evaluated based on the photos taken at every timepoint. To evaluate the degree of cell spreading, the morphology of cells incubated for at least 24-48 h was taken as a reference of complete spreading for each evaluated cell line. Again, the level was categorized in five steps from “completely rounded” to “spreading completed” (0 %, 25 %, 50 %, 75 %, 100 %). The individual values of each triplicate at any time point were then used to generate a heat map based on this five-step categorization.

### Microscopy

Fluorescence microscopy was conducted on a Cell Observer Z1 SD confocal microscope (Zeiss) equipped with 488 nm, 561 nm and 635 nm diode lasers, Plan-Neofluar 10x/0,3, Plan-Neofluar 40x/0,6 and Plan-Apochromat oil 63x/1,4 objectives (Zeiss), a PS1 incubation system (including incubator, heating insert, temp module S and CO_2_ module S, all from Pecon) and an Evolve512 EMCCD camera (Photometrics). Image processing was performed with Zen blue 2.6 (Zeiss) and FIJI 2.1.0.

Live cell microscopy was conducted using 10x or 40x air objectives with the laser intensity as low as possible to prevent bleaching. Usually, images were taken every 7.5 min with Z-stacks containing 6-10 layers at multiple positions for a total of 24-50 h or as indicated in the video captions. Analysis was either performed manually in FIJI or semi-automated using the tools Stardist, TrackMate, and MotilityLab as described below.

Total cell sizes were determined from images taken at 10x objective magnification on a routine light microscope (Zeiss Axiovert 40C) using a OnePlus 8T camera at 12 MP (1,6 µm/px). The image scale was measured using a micrometer calibration slide with defined length units. The cell size was determined using the freehand selection tool in FIJI along the cortex to measure the total area.

### Processing of live microscopy data and single cell tracking

The analysis of time-lapse microscopy data for automated identification of nuclei and tracking of cell movement over time was performed by combining the analysis tools StarDist (Schmidt et al., 2018) (tool for segmentation of cell nuclei in 2D images or stacks based on artificial intelligence), TrackMate (Tinevez et al., 2017) (FIJI plugin to perform single particle tracking of spot-like structures over time) and MotilityLab (http://www.motilitylab.net/) (quantitative and statistical tool to perform cell track analyses).

To prepare microscopy data for the semi-automated process, in each recorded file the nuclear fluorescence channel was extracted, a “Max Intensity”-Z-Projection was performed in FIJI, and the file separately saved in .tif-format. If required for proper target recognition by StarDist, additional contrast enhancement and background subtraction were performed in FIJI.

StarDist is an open-source package for training and implementation of artificial intelligence (AI) approaches to microscopy imaging. First, nuclei was performed. We then trained the AI with either available training datasets (https://zenodo.org/record/3715492#.YdbZQCwxlTa) and training datasets with preset paramters determining the number of iterations, the parameters to improve the recognition of nuclei as well as the progress and validation during the training. Parameters were set as following: number_of_epochs 400, number_of_steps 12, percentage_validation 10, grid_parameter 2, batch_size 4, patch_size 1024, n_rays 32, initial_learning_rate 0.0003.

In a next step, the AI was evaluated and tested on validity and generalizability, again using provided datasets (see link above). A network was described as completely trained once the given curves for “Training loss” and “Validation loss” flattened out. The verified model could then be used to generate predictions from own images. For time-lapse microscopy, the data type was set to “Stacks” representing one image per time point. To allow further processing of generated data in TrackMate, the outputs “Region_of_interest”, “Mask_images”, and “Tracking-file” were generated. The final dataset could then be downloaded as .tif-files and used in TrackMate for further processing. The dataset generated by StarDist consists, among others, of cell tracking files. Each image contains information about the centers of all detected nuclei within a single time point represented as a spot. These spots are recognized by TrackMate and tracked over time. Information on pixel or temporal dimensions are not transferred to tracking files by StarDist and were thus manually entered into the image properties in FIJI. Typical values of data generated by our microscope were a pixel width/height of 1.3333 micron, a voxel depth of 5.00, and a frame interval of 450.00 sec. The tracking file could then be imported to TrackMate with the correct calibration settings. The tracking of individual nuclei was based on the LoG (Laplacian of Gaussian) detector to identify maxima in the images with too close maxima being suppressed. As only the center of individual nuclei was represented in the file, an estimated blob diameter of 1.00 micron and a threshold of 1.00 with sub-pixel localization could be used for cell detection. The tracking of nuclei was based on the LAP (Linear Assignment Problem) algorithm (Jaqaman et al., 2008). Briefly, this algorithm first links spots from frame to frame to generate track segments. Events such as splitting, merging, or gap-closing in case of missing detections are then analyzed in a second step. Typical values for frame linking, gap closing, and segment splitting, or merging were 15.00 micron and a maximum frame gap of 2. The result contained the individual tracks of each cell nucleus over time allowing to save the data as .xml-file for further statistical analysis in MotilityLab or to generate plots using the Chemotaxis and Migration Tool (Ibidi). The online tool MotilityLab (freely available on http://2ptrack.net/plots.php) was used to inspect tracks generated by TrackMate and to exclude possible outliers. Data files from batch analyses can simply be imported and quantified online. We used this platform to organize, rearrange, and extract sub-information from the complete dataset and to arrange data for plotting of the mean square displacement and mean track speed. Further statistical analysis was performed manually using Prism (GraphPad Software).

### Statistical analysis

All statistical analyses were based on datasets of at least three independent experiments (triplicate), if not stated otherwise. Individual datapoints are shown in each graph and where applicable the number of measurements or single values are given in the figure caption. Generally, the mean is presented in each graph and error bars represent the standard deviation (SD) and are shown whenever possible. Details on statistical analyses including parametric/non-parametric, post-hoc test and the multiplicity adjusted P value are given in the respective figure caption. Each dataset was tested for potential outliers using the ROUT (robust regression and outlier removal) method. Briefly, this algorithm is based on the False Discovery Rate (FDR). A predicted model fit is then used to decide whether a data point is far enough away to be considered an outlier. With this method, multiple outliers can be identified and removed simultaneously.

If applicable, datasets were tested for normality and lognormality prior to statistical analysis. Typically, the D’Agostino-Pearson normality test was applied, which first calculates the skewness and kurtosis relative to a Gaussian distribution.

Subsequently, a P value is calculated from the discrepancies of expected and measured values. For P > 0.05, a dataset was defined to follow a normal distribution. If no test for normal distribution was performed and datasets were assumed not to follow a Gaussian distribution, a nonparametric test was conducted, else, either a t test or ANOVA was performed

For distributions that approximately followed a Gaussian distribution, a t test (two groups) or an ANOVA (three or more sets of measurement) was applied. As most datasets were obtained from at least three different groups, commonly a one-way ANOVA (analysis of variance) was performed, followed by an appropriate post-hoc test. Based on these multiple comparisons test, a multiplicity adjusted P value was calculated for each pair. An alpha threshold of 0.05 and confidence level of 95 % has usually been chosen, if not stated differently. Where possible, multiplicity adjusted P values are shown as number in the figure or figure caption. In the interest of legibility, P values were partially represented as asterisks and described in the corresponding figure caption (P Value style GP: 0.1234 (ns), 0.0332 (*), 0.0021 (**), 0.0002 (***), < 0.0001 (****)).

## Supplemementary Material

The following supplementary material is provided:

- Supplementary tables 1-3 in one single pdf file.
- Supplementary figures S1-S5 including figure captions in one single pdf file.
- Videos 1-4 in one zip archive

## Acknowledgements

Wieland Mayer and Matteo Hofmann are acknowledged for technical assistance. The authors sincerely thank Peter Krauss for the donation of the anti-SEPT9 antibody, Ernst-Martin and Annette Füchtbauer for providing the feeder cells and Elias Spiliotis for the EGFP-SEPT9 overexpression plasmid.

Matthias Hecht received financial support by the International Graduate School in Molecular Medicine, Ulm University.

The authors declare no competing financial interests.

## Author contributions

Matthias Hecht performed the experiments, prepared the figures and wrote the manuscript. Pia Marrhofer generated and validated the SEPT9 KO cell line. Nils Johnsson analyzed the data and improved the manuscript. Thomas Gronemeyer analyzed the data, conceived the study and wrote the manuscript.

## Video captions

All videos were imaged by time lapse fluorescence microscopy at 40x magnification 24 h post transfection. Acquisition rate: 1 image every 450 sec. Frame rate: 4 frames/second. Shown is a max intensity Z-projection. Scale bar: 10 µM.

**Video 1.** EPLIN and SEPT9 co-locate along cytosolic septin fibers at the protrusion tip and at the cleavage furrow of dividing cells. Stable GFP-SEPT9 (green) expressing cells were transfected with mRuby-EPLIN (magenta) and stained with SPY-DNA650 (blue) for 15 h before microscopy.

**Video 2:** Visualization of lamellipodia in WT 1306 fibroblasts. Cells were treated with 100 nM SiR Actin (grey) for 15h before microscopy.

**Video 3:** Visualization of lamellipodia in 1306 fibroblasts after EPLIN KD. Cells were transfected with EPLIN siRNA and treated with 100 nM SiR Actin (grey) for 15h before microscopy.

**Video 4:** Visualization of lamellipodia in 1306 fibroblasts transiently overexpressing EPLIN_ΔLIM_. Cells were transfected with GFP-EPLIN_ΔLIM_ and treated with 100 nM SiR Actin (grey) for 15h before microscopy.

## References

Abe, K., and M. Takeichi. 2008. EPLIN mediates linkage of the cadherin-catenin complex to F-actin and stabilizes the circumferential actin belt. Proc. Natl. Acad. Sci. U. S. A. 105:13–19. doi:10.1073/pnas.0710504105.

Beckham, Y., R.J. Vasquez, J. Stricker, K. Sayegh, C. Campillo, and M.L. Gardel. 2014. Arp2/3 inhibition induces amoeboid-like protrusions in MCF10A epithelial cells by reduced cytoskeletal-membrane coupling and focal adhesion assembly. PLoS One. 9. doi:10.1371/JOURNAL.PONE.0100943.

Bowen, J.R., D. Hwang, X. Bai, D. Roy, and E.T. Spiliotis. 2011. Septin GTPases spatially guide microtubule organization and plus end dynamics in polarizing epithelia. J. Cell Biol. doi:10.1083/jcb.201102076.

Calvo, F., R. Ranftl, S. Hooper, A.J. Farrugia, E. Moeendarbary, A. Bruckbauer, F. Batista, G. Charras, and E. Sahai. 2015. Cdc42EP3/BORG2 and Septin Network Enables Mechano-transduction and the Emergence of Cancer-Associated Fibroblasts. Cell Rep. 13:2699–2714. doi:10.1016/J.CELREP.2015.11.052.

Chervin-Pétinot, A., M. Courçon, S. Almagro, A. Nicolas, A. Grichine, D. Grunwald, M.H. Prandini, P. Huber, and D. Gulino-Debrac. 2012. Epithelial Protein Lost In Neoplasm (EPLIN) interacts with α-catenin and actin filaments in endothelial cells and stabilizes vascular capillary network in vitro. J. Biol. Chem. 287:7556–7572. doi:10.1074/JBC.M111.328682.

Chircop, M., V. Oakes, M.E. Graham, M.P.C. Ma, C.M. Smith, P.J. Robinson, and K.K. Khanna. 2009. Cell Cycle The actin-binding and bundling protein, EPLIN, is required for cytokinesis. Cell Cycle. 8:757–64. doi:10.4161/cc.8.5.7878.

Collins, R.J., L.D. Morgan, S. Owen, F. Ruge, W.G. Jiang, and A.J. Sanders. 2018. Mechanistic insights of epithelial protein lost in neoplasm in prostate cancer metastasis. Int. J. Cancer. 143:2537–2550. doi:10.1002/ijc.31786.

Connolly, D., Z. Yang, M. Castaldi, N. Simmons, M.H. Oktay, S. Coniglio, M.J. Fazzari, P. Verdier-Pinard, and C. Montagna. 2011. Septin 9 isoform expression, localization and epigenetic changes during human and mouse breast cancer progression. Breast Cancer Res. 13:R76. doi:10.1186/bcr2924.

Doench, J.G., N. Fusi, M. Sullender, M. Hegde, E.W. Vaimberg, K.F. Donovan, I. Smith, Z. Tothova, C. Wilen, R. Orchard, H.W. Virgin, J. Listgarten, and D.E. Root. 2016. Optimized sgRNA design to maximize activity and minimize off-target effects of CRISPR-Cas9. Nat. Biotechnol. 2015 342. 34:184–191. doi:10.1038/nbt.3437.

Dolat, L., J.L. Hunyara, J.R. Bowen, E.P. Karasmanis, M. Elgawly, V.E. Galkin, and E.T. Spiliotis. 2014. Septins promote stress fiber-mediated maturation of focal adhesions and renal epithelial motility. J. Cell Biol. doi:10.1083/jcb.201405050.

Estey, M.P., C. Di Ciano-Oliveira, C.D. Froese, K.Y.Y. Fung, J.D. Steels, D.W. Litchfield, and W.S. Trimble. 2013. Mitotic Regulation of SEPT9 Protein by Cyclin-dependent Kinase 1 (Cdk1) and Pin1 Protein Is Important for the Completion of Cytokinesis. J. Biol. Chem. 288:30075–30086. doi:10.1074/jbc.M113.474932.

Farrugia, A.J., J. Rodríguez, J.L. Orgaz, M. Lucas, V. Sanz-Moreno, and F. Calvo. 2020. CDC42EP5/BORG3 modulates SEPT9 to promote actomyosin function, migration, and invasion. J. Cell Biol. 219. doi:10.1083/JCB.201912159/VIDEO-4.

Füchtbauer, A., L.B. Lassen, A.B. Jensen, J. Howard, A. De Salas Quiroga, S. Warming, A.B. Sørensen, F.S. Pedersen, and E.-M. Füchtbauer. 2011. Septin9 is involved in septin filament formation and cellular stability*. Biol. Chem. 392:769–777. doi:10.1515/BC.2011.088.

Fuechtbauer, A., L.B. Lassen, A.B. Jensen, J. Howard, A. De Salas Quiroga, S. Warming, A.B. Soerensen, F.S. Pedersen, and E.M. Fuechtbauer. 2011. Septin9 is involved in septin filament formation and cellular stability. Biol. Chem. doi:10.1515/BC.2011.088.

Gardel, M.L., I.C. Schneider, Y. Aratyn-Schaus, and C.M. Waterman. 2010. Mechanical Integration of Actin and Adhesion Dynamics in Cell Migration. http://dx.doi.org/10.1146/annurev.cellbio.011209.122036. 26:315–333. doi:10.1146/ANNUREV.CELLBIO.011209.122036.

Ghosh, I., R.K. Singh, M. Mishra, S. Kapoor, and S.S. Jana. 2021. Switching between blebbing and lamellipodia depends on the degree of non-muscle myosin II activity. J. Cell Sci. 134. doi:10.1242/JCS.248732.

Gilad, R., K. Meir, I. Stein, L. German, E. Pikarsky, and N.J. Mabjeesh. 2015. High SEPT9_i1 Protein Expression Is Associated with High-Grade Prostate Cancers. PLoS One. 10:e0124251. doi:10.1371/journal.pone.0124251.

Gulino-Debrac, D. 2013. Mechanotransduction at the basis of endothelial barrier function. Tissue Barriers. 1:e24180. doi:10.4161/tisb.24180.

Han, M.-Y., H. Kosako, T. Watanabe, and S. Hattori. 2007. Extracellular Signal-Regulated Kinase/Mitogen-Activated Protein Kinase Regulates Actin Organization and Cell Motility by Phosphorylating the Actin Cross-Linking Protein EPLIN. Mol. Cell. Biol. 27:8190–8204. doi:10.1128/MCB.00661-07/SUPPL_FILE/VIDEO4.ZIP.

Hecht, M., R. Rösler, S. Wiese, N. Johnsson, and T. Gronemeyer. 2019. An Interaction Network of the Human SEPT9 Established by Quantitative Mass Spectrometry. G3 Genes|Genomes|Genetics. g3.400197.2019. doi:10.1534/g3.119.400197.

Janiszewska, M., M.C. Primi, and T. Izard. 2020. Cell adhesion in cancer: Beyond the migration of single cells. J. Biol. Chem. 295:2495. doi:10.1074/JBC.REV119.007759.

Jaqaman, K., D. Loerke, M. Mettlen, H. Kuwata, S. Grinstein, S.L. Schmid, and G. Danuser. 2008. Robust single-particle tracking in live-cell time-lapse sequences. Nat. Methods 2008 58. 5:695–702. doi:10.1038/nmeth.1237.

Joo, E.E., and K.M. Yamada. 2014. MYPT1 regulates contractility and microtubule acetylation to modulate integrin adhesions and matrix assembly. Nat. Commun. 2014 51. 5:1–13. doi:10.1038/ncomms4510.

Kasai, N., A. Kadeer, M. Kajita, S. Saitoh, S. Ishikawa, T. Maruyama, and Y. Fujita. 2018. The paxillin-plectin-EPLIN complex promotes apical elimination of RasV12-transformed cells by modulating HDAC6-regulated tubulin acetylation. Sci. Rep. 8. doi:10.1038/S41598-018-20146-1.

Lam, M., and F. Calvo. 2019. Regulation of mechanotransduction: Emerging roles for septins. Cytoskeleton. 76:115–122. doi:10.1002/CM.21485.

Linklater, E.S., E.D. Duncan, K.-J. Han, A. Kaupinis, M. Valius, T.R. Lyons, and R. Prekeris. 2021. Rab40/Cullin5 complex regulates EPLIN and actin cytoskeleton dynamics during cell migration and invasion. biorxiv. doi:10.1101/2021.04.01.438077.

Longtine, M.S., D.J. DeMarini, M.L. Valencik, O.S. Al-Awar, H. Fares, C. De Virgilio, and J.R. Pringle. 1996. The septins: roles in cytokinesis and other processes. Curr. Opin. Cell Biol. 8:106–19. doi:10.1016/s0955-0674(96)80054-8.

Marcus, J., M. Bejerano-Sagie, N. Patterson, S. Bagchi, V. V. Verkhusha, D. Connolly, G.L. Goldberg, A. Golden, V.P. Sharma, J. Condeelis, and C. Montagna. 2019. Septin 9 isoforms promote tumorigenesis in mammary epithelial cells by increasing migration and ECM degradation through metalloproteinase secretion at focal adhesions. Oncogene. 38:5839–5859. doi:10.1038/s41388-019-0844-0.

Maul, R.S., and D.D. Chang. 1999. EPLIN, Epithelial protein lost in neoplasm. Oncogene. 18:7838–7841. doi:10.1038/sj.onc.1203206.

Maul, R.S., Y. Song, K.J. Amann, S.C. Gerbin, T.D. Pollard, and D.D. Chang. 2003. EPLIN regulates actin dynamics by cross-linking and stabilizing filaments. J. Cell Biol. 160:399–407. doi:10.1083/jcb.200212057.

Mavrakis, M., Y. Azou-Gros, F.-C. Tsai, J. Alvarado, A. Bertin, F. Iv, A. Kress, S. Brasselet, G.H. Koenderink, and T. Lecuit. 2014. Septins promote F-actin ring formation by crosslinking actin filaments into curved bundles. Nat. Cell Biol. 16:322–334. doi:10.1038/ncb2921.

McMurray, M., and J. Thorner. 2019. Turning it inside out: The organization of human septin heterooligomers. Cytoskeleton (Hoboken). 76:449–456. doi:10.1002/CM.21571.

Mendonça, D.C., J.N. Macedo, S.L. Guimarães, F.L. Barroso da Silva, A. Cassago, R.C. Garratt, R. V. Portugal, and A.P.U. Araujo. 2019. A revised order of subunits in mammalian septin complexes. Cytoskeleton. 76:457–466. doi:10.1002/cm.21569.

Michelsen, J.W., K.L. Schmeichel, M.C. Beckerle, and D.R. Winge. 1993. The LIM motif defines a specific zinc-binding protein domain. Proc. Natl. Acad. Sci. U. S. A. 90:4404–4408. doi:10.1073/PNAS.90.10.4404.

Ohashi, T., M. Idogawa, Y. Sasaki, and T. Tokino. 2017. p53 mediates the suppression of cancer cell invasion by inducing LIMA1/EPLIN. Cancer Lett. 390:58–66. doi:10.1016/j.canlet.2016.12.034.

Pal, M., S. Bhattacharya, G. Kalyan, and S. Hazra. 2018. Cadherin profiling for therapeutic interventions in Epithelial Mesenchymal Transition (EMT) and tumorigenesis. Exp. Cell Res. 368:137–146. doi:10.1016/j.yexcr.2018.04.014.

Ran, F.A., P.D. Hsu, J. Wright, V. Agarwala, D.A. Scott, and F. Zhang. 2013. Genome engineering using the CRISPR-Cas9 system. Nat. Protoc. 8:2281–2308. doi:10.1038/nprot.2013.143.

Schmidt, U., M. Weigert, C. Broaddus, and G. Myers. 2018. Cell Detection with Star-convex Polygons. Lect. Notes Comput. Sci. (including Subser. Lect. Notes Artif. Intell. Lect. Notes Bioinformatics). 11071 LNCS:265–273. doi:10.1007/978-3-030-00934-2_30.

Shindo, A., A. Audrey, M. Takagishi, M. Takahashi, J. Wallingford, and M. Kinoshita. 2018. Septin-dependent remodeling of cortical microtubule drives cell reshaping during epithelial wound healing. J. Cell Sci. 131. doi:10.1242/JCS.212647.

Shuman, B., and M. Momany. 2022. Septins From Protists to People. Front. cell Dev. Biol. 9. doi:10.3389/FCELL.2021.824850.

Simi, A.K., A.A. Anlaş, M. Stallings-Mann, S. Zhang, T. Hsia, M. Cichon, D.C. Radisky, and C.M. Nelson. 2018. A Soft Microenvironment Protects from Failure of Midbody Abscission and Multinucleation Downstream of the EMT-Promoting Transcription Factor Snail. Cancer Res. 78:2277–2289. doi:10.1158/0008-5472.CAN-17-2899.

Smith, C., L. Dolat, D. Angelis, E. Forgacs, E.T. Spiliotis, and V.E. Galkin. 2015. Septin 9 Exhibits Polymorphic Binding to F-Actin and Inhibits Myosin and Cofilin Activity. J. Mol. Biol. 427:3273–3284. doi:10.1016/j.jmb.2015.07.026.

Song, L., and Y. Li. 2015. SEPT9: : A Specific Circulating Biomarker for Colorectal Cancer. Adv. Clin. Chem. 72:171–204. doi:10.1016/bs.acc.2015.07.004.

Soroor, F., M.S. Kim, O. Palander, Y. Balachandran, R.F. Collins, S. Benlekbir, J. Rubinstein, and W.S. Trimble. 2020. Revised subunit order of mammalian septin complexes explains their in-vitro polymerization properties. Mol. Biol. Cell. mbc.E20-06-0398. doi:10.1091/mbc.E20-06-0398.

Sun, B.O., Y. Fang, Z. Li, Z. Chen, and J. Xiang. 2015. Role of cellular cytoskeleton in epithelial-mesenchymal transition process during cancer progression. Biomed. reports. 3:603–610. doi:10.3892/br.2015.494.

Surka, M.C., C.W. Tsang, and W.S. Trimble. 2002. The mammalian septin MSF localizes with microtubules and is required for completion of cytokinesis. Mol. Biol. Cell. 13:3532–45. doi:10.1091/mbc.E02-01-0042.

Taguchi, K., T. Ishiuchi, and M. Takeichi. 2011. Mechanosensitive EPLIN-dependent remodeling of adherens junctions regulates epithelial reshaping. J. Cell Biol. 194:643–656. doi:10.1083/JCB.201104124/VIDEO-10.

Taha, M., M. Aldirawi, S. März, J. Seebach, M. Odenthal-Schnittler, O. Bondareva, V. Bojovic, T. Schmandra, B. Wirth, M. Mietkowska, K. Rottner, and H. Schnittler. 2019. EPLIN-α and -β Isoforms Modulate Endothelial Cell Dynamics through a Spatiotemporally Differentiated Interaction with Actin. Cell Rep. 29:1010–1026.e6. doi:10.1016/J.CELREP.2019.09.043.

Tinevez, J.Y., N. Perry, J. Schindelin, G.M. Hoopes, G.D. Reynolds, E. Laplantine, S.Y. Bednarek, S.L. Shorte, and K.W. Eliceiri. 2017. TrackMate: An open and extensible platform for single-particle tracking. Methods. 115:80–90. doi:10.1016/J.YMETH.2016.09.016.

Tokhtaeva, E., J. Capri, E.A. Marcus, J.P. Whitelegge, V. Khuzakhmetova, E. Bukharaeva, N. Deiss-Yehiely, L.A. Dada, G. Sachs, E. Fernandez-Salas, and O. Vagin. 2015. Septin dynamics are essential for exocytosis. J. Biol. Chem. doi:10.1074/jbc.M114.616201.

Verdier-Pinard, P., D. Salaun, H. Bouguenina, S. Shimada, M. Pophillat, S. Audebert, E. Agavnian, S. Coslet, E. Charafe-Jauffret, T. Tachibana, and A. Badache. 2017. Septin 9_i2 is downregulated in tumors, impairs cancer cell migration and alters subnuclear actin filaments. Sci. Rep. 7:44976. doi:10.1038/srep44976.

Xu, D., A. Liu, X. Wang, Y. Chen, Y. Shen, Z. Tan, and M. Qiu. 2018. Repression of Septin9 and Septin2 suppresses tumor growth of human glioblastoma cells. Cell Death Dis. 9. doi:10.1038/S41419-018-0547-4.

Yeh, Y.-T., S.S. Hur, J. Chang, K.-C. Wang, J.-J. Chiu, Y.-S. Li, and S. Chien. 2012. Matrix Stiffness Regulates Endothelial Cell Proliferation through Septin 9. PLoS One. 7:e46889. doi:10.1371/JOURNAL.PONE.0046889.

Zeng, Y., Y. Cao, L. Liu, J. Zhao, T. Zhang, L. Xiao, M. Jia, Q. Tian, H. Yu, S. Chen, and Y. Cai. 2019. SEPT9_i1 regulates human breast cancer cell motility through cytoskeletal and RhoA/FAK signaling pathway regulation. Cell Death Dis. 10. doi:10.1038/s41419-019-1947-9.

Zhang, S., X. Wang, A.O. Osunkoya, S. Iqbal, Y. Wang, Z. Chen, S. Müller, Z. Chen, S. Josson, I.M. Coleman, P.S. Nelson, Y.A. Wang, R. Wang, D.M. Shin, F.F. Marshall, O. Kucuk, L.W.K. Chung, H.E. Zhau, and D. Wu. 2011. EPLIN downregulation promotes epithelial-mesenchymal transition in prostate cancer cells and correlates with clinical lymph node metastasis. Oncogene. 30:4941–4952. doi:10.1038/onc.2011.199.

