## Supplementary Figures for "The concerted action of SEPT9 and EPLIN modulates the adhesion and migration of human fibroblasts"

### Supplementary Figure 1

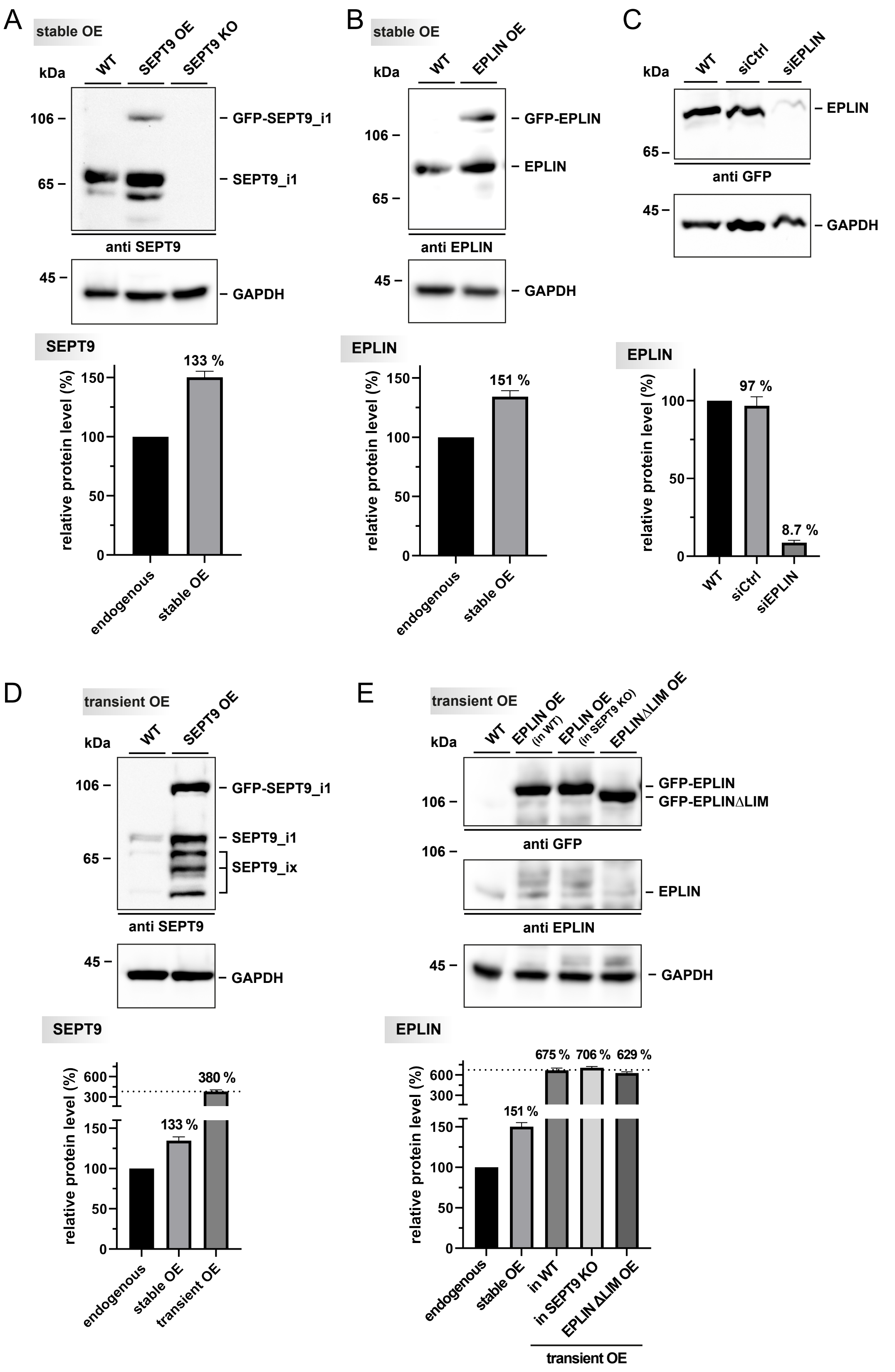

**Figure S1.** Characterization of SEPT9 and EPLIN OE cells as well as EPLIN siRNA mediated KD. **(A)** Western blot and relative quantification of the OE and CRISPR/Cas9-mediated KO of SEPT9. The quantification upon stable OE (1.33-fold) was based on the endogenous level of SEPT9 in WT cells. **(B)** Western blot and relative quantification of the stable OE of EPLIN indicate a 1.5-fold overexpression compared to the WT. **(C)** Western blot and relative quantification of 1306 cells treated either with non-targeting siRNA or siRNA against EPLIN for 48 h. **(D)** Western blot and relative quantification of transient OE of GFP-SEPT9. **(E)** Western blot and relative quantification of transient OE of GFP-EPLIN in WT fibroblasts and in SEPT9 KO cells as well as transient OE of GFP-EPLIN$_{\square \mathrm{LIM}}$.

The quantitative data are based on triplicate analysis and depict the means ± SD.

#### Supplementary Figure 2

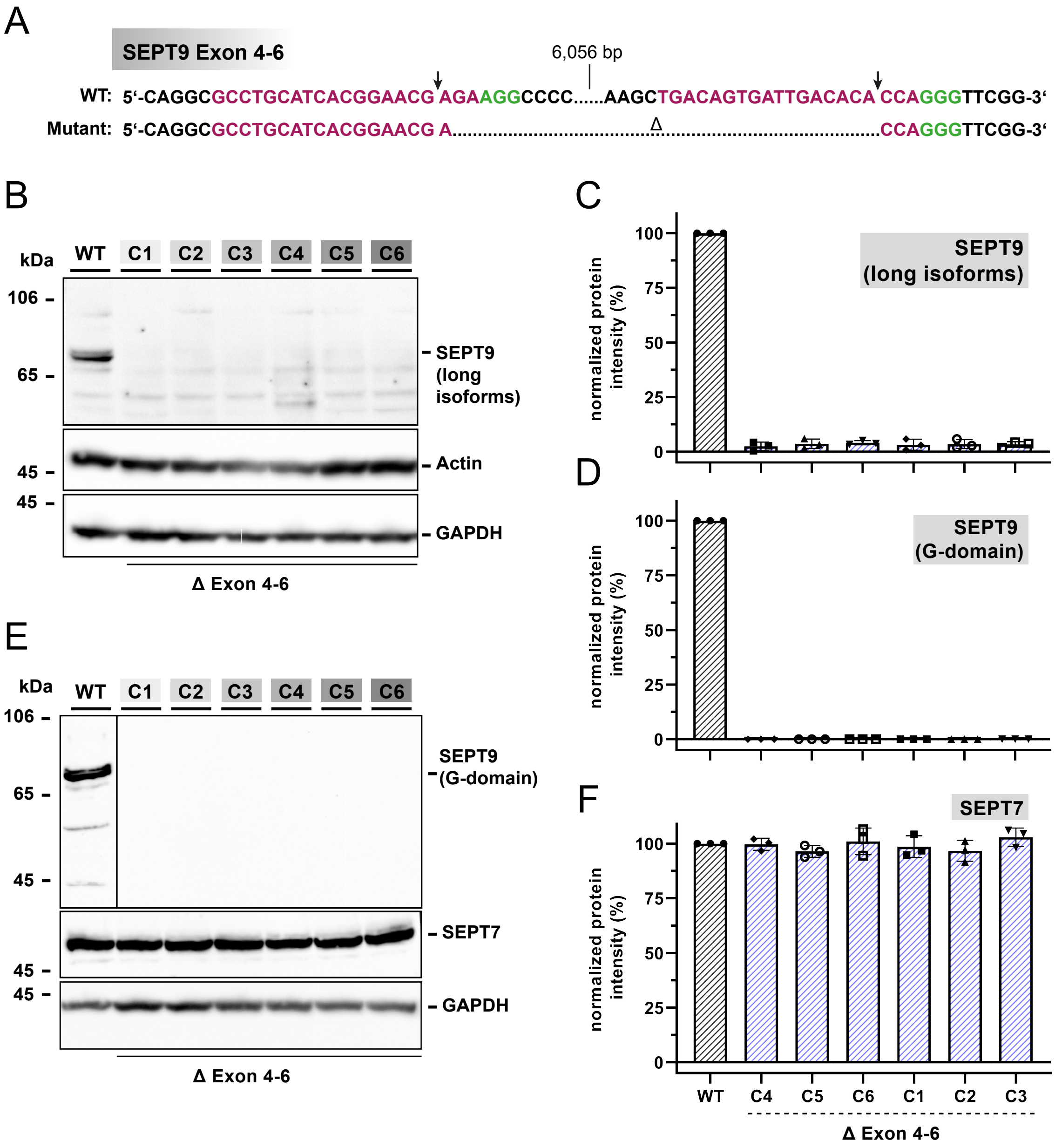

**Figure S2.** Characterization of the SEPT9 KO cell line. **(A)** Restriction sites for Cas9 5' of exon 4 and 3' of exon 6 (labelled with an arrow) of the SEPT9 gene. The PAM sites are indicated by green letters. **(B, C)** The western blot from 1306 cell lysates using an antibody covering all long isoforms of SEPT9 confirmed the absence of a SEPT9 expression product in in all tested clones without affecting the actin expression. The quantification is shown relative to the WT level. **(D, E, F)** The western blot against the SEPT9 G-domain confirmed the depletion of SEPT9 in in all tested clones. Expression of SEPT7 was not altered. All Western blots were performed with a sample size of at least n=3. All quantitative data are depicted as means ± SD. C1-C6 indicate different clones of the Δexon4-6 KO cell line.

#### Supplementary Figure 3

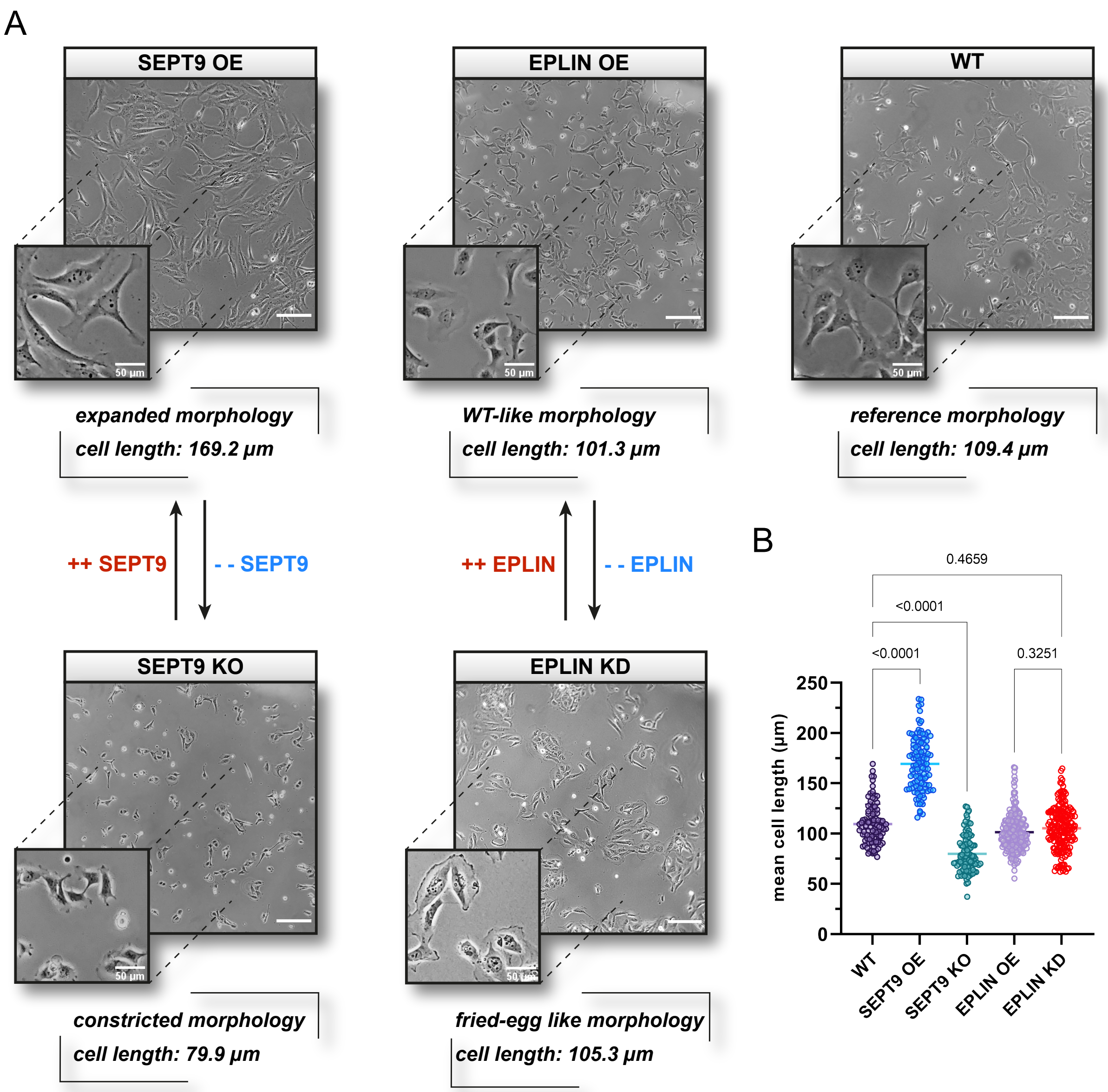

**Figure S3.** Morphological influence of SEPT9 and EPLIN in 1306 fibroblast cells. **(A)** 1306 fibroblasts with an elevated level of SEPT9 showed an expanded morphology whereas a knock-out induced a severely constricted phenotype. The overexpression of EPLIN had no significant influence on the overall morphology, whereas the cell-matrix contact area increased with a fried-egg like morphology in EPLIN KD cells. EPLIN KD cell were treated with siRNA 48 h before documentation, all other cell lines were stably mutated. **(B)** Quantification of the mean cell length. Upon overexpression of SEPT9 the length of 1306 cells increased to approximately 150 %, with a decrease in cell length to roughly 70 % upon SEPT9 KO compared to WT cells. Despite a change in the morphology of EPLIN KO cells, no measurable changes in cell length were determined upon OE or KO of EPLIN. Light microscopy images were taken with a 10x magnification objective (scale bar = 200 µm). All quantitative data were collected from three independent experiments with n=33 per replicate and are depicted as means ± SD. Significance values were calculated by one-way ANOVA followed by Tukey's multiple comparison test.

### Supplementary Figure 4

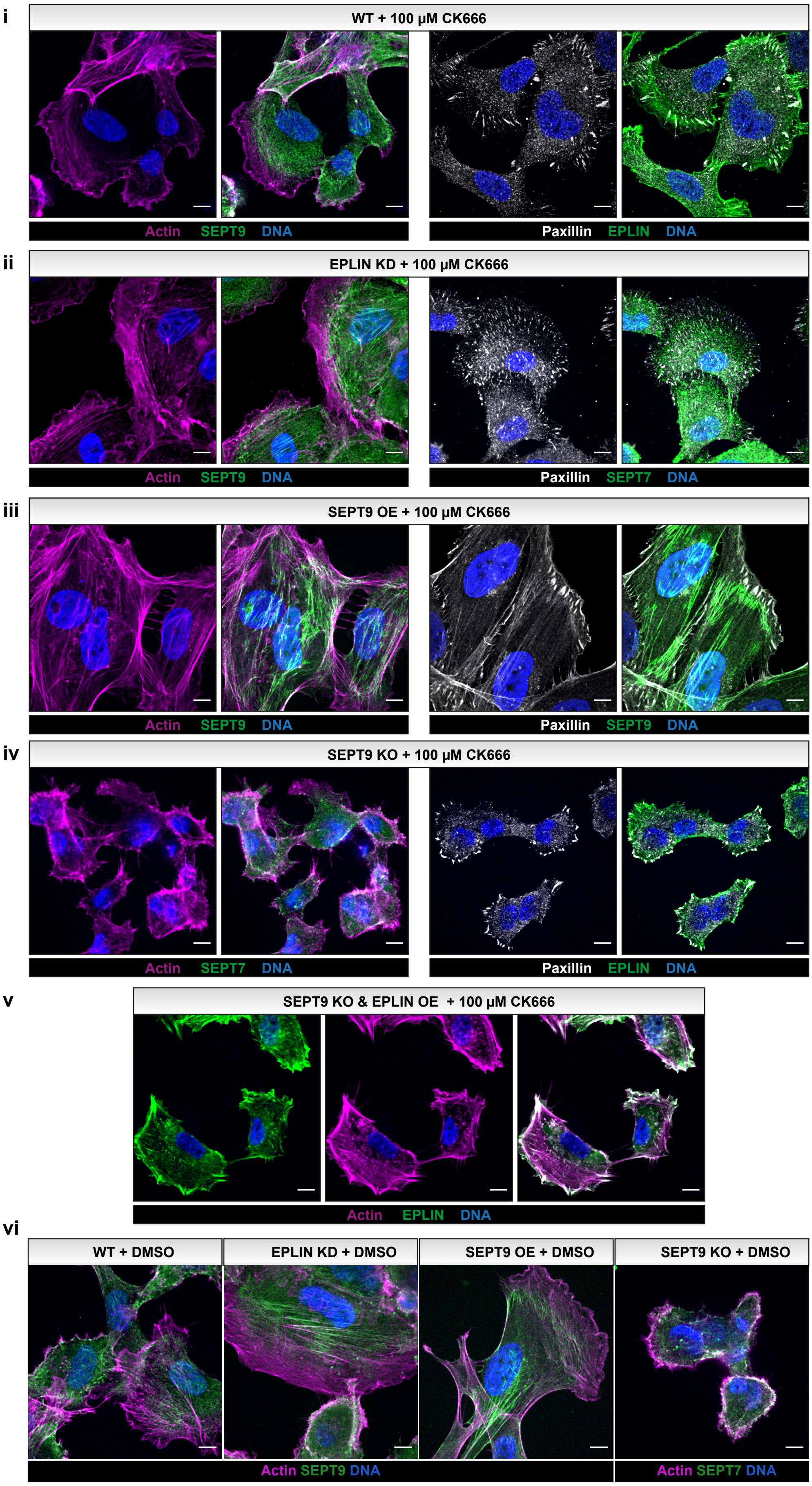

**Figure S4.** 1306 cells with SEPT9 OE or EPLIN KD maintain stabilized actin structures upon treatment with CK666. Immunostained (i-v) and GFP-labelled SEPT9 (ii) or EPLIN (v) in 1306 fibroblasts upon treatment with 100 μM CK666 for 30 min (nuclei were stained with SPY650-DNA dye). All cells except SEPT9 KO cells (iv) showed an actin network enriched in transverse arcs. EPLIN KD cells had a SEPT9 OE-like stabilized cortical actin network compared to the WT, even upon CK666 treatment. Alike non-treated cells, focal adhesions were not restricted to the plasma membrane. (scale bar = 10 μM)

**Supplementary Figure 5**

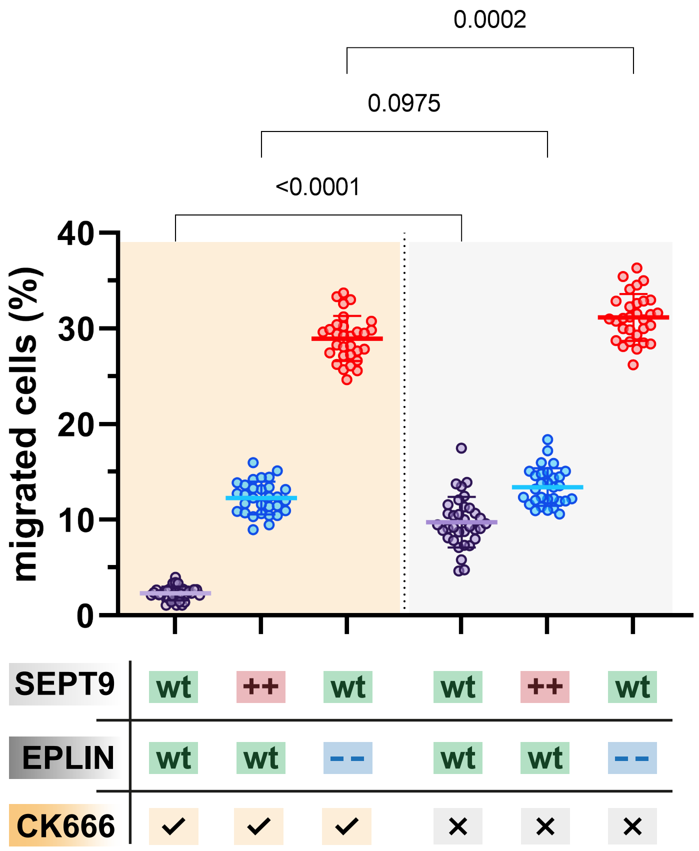


**Figure S5.** 1306 fibroblasts are insensitive to CK666 treatment upon SEPT9 OE or EPLIN KD. The cell migration recorded in a transwell assay was significantly reduced in WT cells upon treatment with 100 µM of the Arp2/3 inhibitor CK666 but only slightly upon SEPT9 OE or EPLIN KD.
