## Supplementary Tables for "The concerted action of SEPT9 and EPLIN modulates the adhesion and migration of human fibroblasts"

### Supplementary Information

***Suppl. Table 1:*** *Oligonucleotides as primers for PCRs (in 5'-3 orientation)*

| Name | Sequence (5'-3') | Construct/Purpose |
|---|---|---|
| GSTEPLa_SbfI_fw | GATCCCTGCAGGCTGAAAATTGTCTAGGAGAATCCAGGC | GST-EPLIN |
| GSTEPLa_AscI_rv | CGATGGCGCGCCTCACTCTTCATCCTCATCCT | GST-EPLIN |
| EpLIM_BamHI_fw | GTCAGGATCCGAGACCTGCGTGGAATGTCAG | GST-EPLIN_LIM |
| EplLIM_AscI_rv | GTAGGCGCGCCTCAAGATTTAAAGAGTTGATTGAAGTGA GGC | GST-EPLIN_LIM |
| Sept9-Sfi1for | CCGAAGGCCAGCACGGCCGAAAACCTGTACTTCCAGGGT AAGAAGTCTTACTCAGGAGG | GST-SEPT9, His-SEPT9 |
| Sept9-Sfi1rev | CTGTGGGCCAAAAAGGCCTTATCACATCTCTGGGGCTTCTGG | GST-SEPT9, His-SEPT9 |
| His-EPLIN_Sfi_fw | GCATCGGCCAGCACGGCCGAAAATTGTCTAGGAGAATC CAGGC | His-EPLIN |
| His-EPLIN_Sfi_rv | GTACTGGCCAAAAAGGCCTCACTCTTCATCCTCATCCTC ATCA | His-EPLIN |
| GFP-EPLIN_SalI_fw | CTAGGTCGACATGGTGAGCAAGGGCGAGG | EGFP-EPLIN |
| GFP-EPLIN_NheI_rv | CTAGGCTAGCTCACTCTTCATCCTCATCCTCATCATAATA C | EGFP-EPLIN |
| iRFP_Crispr.rev | GACTGAATTCTTAGCGTTGGTGGTGGGC | pSpCas9(BB)-2A-iRFP |
| T2A_iRFP-fw | GACTGAATTCGGCAGTGGAGAGGGCAG | pSpCas9(BB)-2A-iRFP |
| SEPT9_X4seq.fw | GTTCCGCTCTAACTCCTCTGC | sequencing of SEPT9 Exon4 |
| SEPT9_X4seq.rv | GCTGGTTTCCGGCATTGG | sequencing of SEPT9 Exon4 |
| SEPT9_X6seq.fw | GTGCAGATATTGAGGAGAAAGGCG | sequencing of SEPT9 Exon6 |
| SEPT9_X6seq.rv | GGCCTCACCAGTTCTCGTTG | sequencing of SEPT9 Exon6 |
| S9_Gdom_sfi_fw | GCATCGGCCAGCACGGCCCAGGGCTTCGAGTTCAACAT CATG | His-SEPT9 ΔNΔC |
| S9_Gdom_sfi_rv | CATGAGGCCAAAAAGGCCTCACTCGTTGAGGCGCTTCA CAC | His-SEPT9 ΔNΔC |
| S9_Sfi_fw | CCATGAGCAGCCATCATCATC | His-SEPT9 ΔC |
| S9_sfi_rv | GTTAGCAGCCGGATCCGTTG | His-SEPT9 ΔN |
| Primer 1_EPLIN_splic1.fw | CATGGACGAGCTGTACAAGTCC | EGFP-EPLIN ΔLIM |
| Primer 2_EPLIN_splic2.rv | CATCATAGTTGCCCTTTCTTGCAGGTGCCTGAAAC | EGFP-EPLIN ΔLIM |
| Primer 3_EPLIN_splic3.fw | CAGGCACCTGCAAGAAAGGGCAACTATGATGAAGGC | EGFP-EPLIN ΔLIM |
| Primer 4_EPLIN_splic4.rv | CTAGATCCGGTGGATCCCGG | EGFP-EPLIN ΔLIM |
| GFP-LIM_BspEI_fw | GTCATCCGGAGAGACCTGCGTGGAATGTCAGAAG | EGFP-EPLIN_LIM |
| GFP-LIM_SalI_rv | CATGGTCGACATCAAGATTTAAAGAGTTGATTGAAGTGA GGC | EGFP-EPLIN_LIM |

**Suppl. Table 2: Primary and secondary antibodies (Abs) used for immunofluorescence (IF) or Western Blotting (WB)**

| Product | Host | Dilution IF | Dilution WB | Supplier | Product ID |
|---|---|---|---|---|---|
| **Primary Abs** | | | | | |
| α-SEPT9 (N-term) | rabbit | 1:100 | 1:5000 | Prof. Krauss, Leibniz Institute of Molecular Pharmacology, Berlin | |
| α-SEPT9 | mouse | - | 1:3000 | Sigma Aldrich (2C6) | # WH001-0801M1 |
| α-SEPT9 (G-domain) | rabbit | 1:100 | 1:5000 | Bethyl Laboratories | # A302-354A |
| α-SEPT7 | rabbit | 1:100 | 1:5000 | Sigma-Aldrich | # HPA029524 |
| α-β-Actin | mouse | 1:100 | 1:2000 | SCBT (E-10) | # sc-365791 |
| α-EPLIN | rabbit | 1:200-1:100 | - | FineTest | # FNab02812 |
| α-EPLIN | mouse | 1:100 | 1:2500 | SCBT (20) | # sc-136399 |
| α-Paxillin | mouse | 1:100 | 1:3000 | SCBT (D-9) | # sc-365174 |
| α-GAPDH-HRP | mouse | 1:100 | 1:2500 | SCBT (0411) | # sc-47724 |
| α-$His_6$ | mouse | - | 1:5000 | Sigma Aldrich (HIS-1) | # H1029 |
| **Secondary Abs** | | | | | |
| α-mouse-Alexa Fluor® 488 | goat | 1:1000-1:500 | - | Thermo Fisher Scientific | # A28175 |
| α-mouse-Alexa Fluo® 555 | goat | 1:1000-1:500 | - | Thermo Fisher Scientific | # A32727 |
| α-mouse-Alexa Fluor® 647 | goat | 1:1000-1:500 | - | Thermo Fisher Scientific | # A32728 |
| α-rabbit-Alexa Fluor® 488 | goat | 1:1000-1:500 | - | Thermo Fisher Scientific | # A27034 |
| α-rabbit-Alexa Fluor® 555 | goat | 1:1000-1:500 | - | Thermo Fisher Scientific | # A27039 |
| α-mouse-HRP | goat | - | 1:5000 | Sigma Aldrich | # A4416 |
| α-rabbit-HRP | goat | - | 1:5000 | Bio-Rad | # 170-6515 |

***Suppl. Table 3: Sequences of sgRNAs for CRISPR/Cas9 gene knockout*** *(capital letters mark the targeting sgRNA sequence, whereas lowercase letters indicate overhangs for cloning into pSpCas9(BB)2A via BbsI.*

| Name | Sequence (5'-3') | PAM site | On target value [%] | Off-target value [%] |
|---|---|---|---|---|
| **SEPT9_sgRNA_Exon 4B_fw** | caccgGCCTGCATCACGGAACGAGA | AGG | 62.5 | 90.3 |
| **SEPT9_sgRNA_Exon 4B_rv** | aaacTCTCGTTCCGTGATGCAGGCc | | | |
| SEPT9_sgRNA_Exon 6E_fw | caccgTGACAGTGATTGACACACCA | CGG | 76.7 | 66.4 |
| SEPT9_sgRNA_Exon 6E_rv | aaacTGGTGTGTCAATCACTGTCAc | | | |
